# Transient MYC Mimicking the Exercise Response Orchestrates Multifaceted Skeletal Muscle Adaptations

**DOI:** 10.64898/2026.06.29.735332

**Authors:** Ronald G. Jones, Pieter J. Koopmans, Nathan Serrano, Ana Regina Cabrera, Francielly Morena, Toby L. Chambers, Ruth Walters, Agnieszka K. Borowik, Michael E. Taylor, Ross P. Wohlgemuth, Jazmine A. Eccles Miller, Abigail L. Zirbel, Lea C. Range, Sebastian Edman, Ferdinand von Walden, Christopher S. Fry, Joanna L. Fiddler, Benjamin F. Miller, Ahmed Ismaeel, Yuan Wen, Kevin A. Murach

## Abstract

Despite decades of study, MYC’s physiological role in adult tissues remains obscured by the models used to investigate it. Most work relies almost exclusively on chronic or constitutive MYC induction that recapitulates the sustained activity found in tumorigenesis and expectedly produces pathological outcomes. What has gone largely untested is how MYC operates when induced in a controlled and physiologically relevant manner, as it is during adaptive processes such as exercise. Using a recombination-independent strategy in adult skeletal muscle, transient MYC bursts drive coordinated hypertrophic-metabolic reprogramming followed by a shift in myosin fiber type, recapitulating adaptations characteristic of concurrent endurance and resistance training. A single pleiotropic transcription factor governing several aspects of muscle health reframes perspectives on a gene understood almost entirely in the context of pathology. We uncover a previously unrecognized muscle-specific role for MYC in controlling muscle cell composition and present a multi-timepoint multi-omic resource for interrogating MYC: data.myoanalytics.com/study/myc_transient/gene.

## Introduction

*c-MYC* is among the most studied genes in mammalian biology. Dysregulated in an estimated 70% of human cancers ^1^, *MYC*/*Myc* encodes a transcription factor (MYC) that drives proliferative cell growth across virtually all cell types ^2,3^. Despite decades of investigation, however, the physiological role of MYC in differentiated adult tissues remains poorly understood. This gap is not for lack of effort but rather a consequence of the experimental models used to study MYC. *In vivo* work relies almost exclusively on chronic or constitutive MYC induction; approaches that, by design, recapitulate the sustained and unregulated activity characteristic of pathology ^4^. Under these persistent conditions, deleterious outcomes from MYC are not surprising but predetermined by the model ^5^. What has gone largely untested is how MYC functions when expressed on a physiological timescale *in vivo*: transiently induced then rapidly degraded, as occurs during normal adaptive processes in adult tissues.

The distinction between the effects of acute and chronic induction of a transcription factor is not merely quantitative. In transcription factor biology, the temporal pattern of activation can determine qualitatively different cellular outcomes ^6^. One salient example is what happens with *in vivo* expression of the Yamanaka reprogramming factors OCT3/4, SOX2, KLF4, and MYC (OSKM) ^7^. Consistent expression causes teratomas ^8^ whereas cyclic short-term induction of the same factors can ameliorate hallmarks of aging and extend lifespan without oncogenic transformation ^9^. MYC is a component of this reprogramming cassette and among the most tightly regulated and rapidly turned over transcription factors in the cell ^10–12^. MYC has architectural and functional features at the gene locus that actively suppress spurious expression ^13–15^. The stringency of endogenous MYC regulation suggests that the duration and magnitude of its induction are intrinsic to its normal functioning ^11^; and yet, the vast majority of experimental studies override these precise regulatory parameters ^5^.

Skeletal muscle provides a uniquely informative system to explore the function of MYC. As the largest organ by mass, its most voluminous cells (muscle fibers) are a post-mitotic syncytium that are resistant to primary tumor formation ^16,17^; however, MYC transcript and protein is prominently upregulated in muscle fiber nuclei (myonuclei) in response to stress ^18–22^. MYC is transiently enriched in muscle following both resistance and endurance exercise in humans, peaking within hours and returning to baseline within a day ^23^. It is the most influential transcription factor governing the late-stage transcriptional response to a bout of resistance exercise in human skeletal muscle ^24^, and is the Yamanaka factor most induced by exercise ^25,26^. As expected, chronic induction of MYC in murine skeletal muscle is deleterious for muscle health and function but does not cause oncogenesis ^4,5^. Short-term induction of the full OSKM cassette (including MYC) specifically in myofibers promotes muscle regeneration by remodeling the stem cell niche ^27^. These observations reinforce skeletal muscle as a tissue in which controlled transient transcription factor activity drives adaptive rather than pathological outcomes. In the aforementioned contexts, MYC operates in cells that do not proliferate, removing the dominant confound of cancer-prone cell types and models.

Here, we use a skeletal muscle fiber-specific recombination-independent non-constitutive doxycycline-inducible system (called HSA-MYC) to deliver controlled transient bursts of MYC in adult mice. We hypothesized that repeated transient MYC over an extended time course would result in hypertrophy that is dominantly defined by ribosome biogenesis-related processes across omic layers ^23,28–31^. Instead, we reveal that pulsatile MYC is sufficient to drive oxidative and biosynthesis-oriented metabolic changes that culminate in a myosin heavy chain (MyHC) fiber type transition, concomitant with muscle hypertrophy. We therefore expose MYC as a single exercise-induced transcription factor that coordinates several aspects of muscle health via repeated short bursts. The MYC-conditioned muscle phenotype captures cellular, metabolic, and molecular hallmarks of combined endurance+resistance exercise training ^32–34^. Multi-timepoint transcriptomic, proteomic, and single-nucleus RNA-sequencing data define the temporal architecture of MYC-mediated adaptation. These data are provided as an open interactive resource for the skeletal muscle and MYC biology communities.

No longer obscured by the very models used to investigate it, we recast MYC as a gene with potent and multifaceted functions in a differentiated tissue that are only demystified by replicating its natural physiological cadence. Our approach reveals MYC’s previously unrecognized muscle-specific role in controlling the proportional composition of contractile proteins pursuant to a shift in metabolism. The data we provide are a platform for understanding the effects of transient MYC in a non-proliferative cancer-resistant cell type. This information could open new possibilities for harnessing the powerful effects of MYC via canonical as well as unique downstream targets in skeletal muscle.

## Results

### A single transient pulse of MYC in murine soleus muscle causes transcriptional changes related to translational enhancement, metabolism, and muscle identity

To define the acute transcriptional consequences of myofiber-restricted MYC induction, we delivered doxycycline in drinking water to young adult female HSA-MYC mice for 48 hours and harvested soleus muscle 12 hours following doxycycline removal (Figure 1A). We previously established that 12 hours after the removal of doxycycline in drinking water is when MYC protein is elevated in skeletal muscle ^25^. MYC protein returns to baseline levels within 24 hours of doxycycline removal in our model ^24,25^. Bulk RNA sequencing was performed on soleus muscle from MYC-induced (HSA-MYC) and control doxycycline-treated littermate HSA-only mice (n=3/group). The soleus was chosen due to the hypertrophy we observe in this muscle with MYC pulses in our previous work ^24^, but also because the soleus is the only murine hindlimb skeletal muscle with a MyHC fiber type proportion analogous to most human leg muscles: approximately equal parts MyHC I and IIa without pure IIx or any IIb ^35,36^. Using the soleus adds a translational aspect to our analyses. Principal component analysis (PCA) confirmed a large separation between groups: PC1 and PC2 explain 42.8% and 20.3% of total variance, respectively. MYC samples clustered tightly and distinctly from controls along the primary axis of variance (Figure 1B). This degree of separation indicates that myofiber-restricted MYC imposes a dominant and coherent transcriptional shift, consistent with MYC’s role as a powerful transcription factor that influences an appreciable proportion of the genome ^37^.

**Figure 1.**
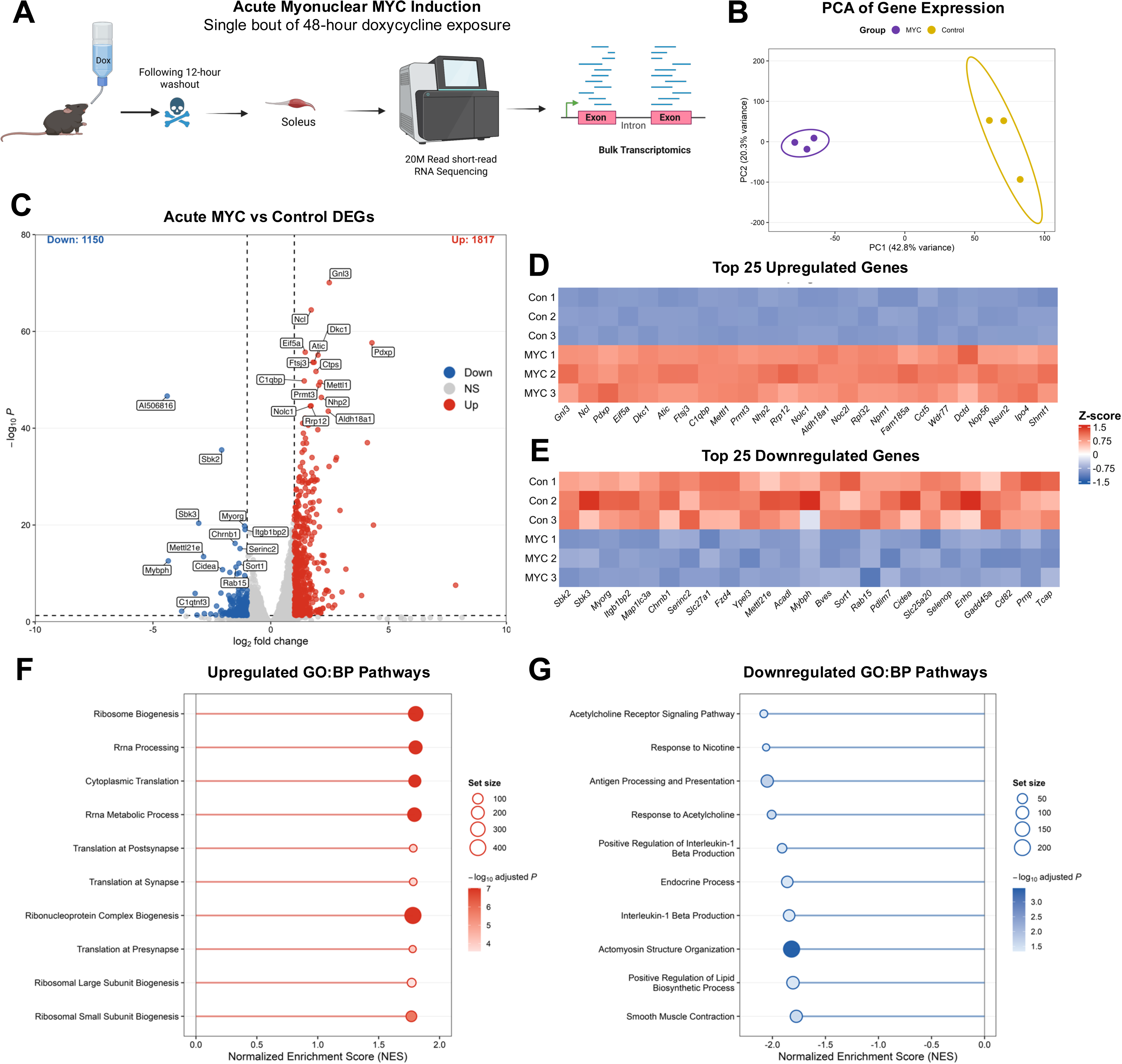
Acute MYC induction rapidly remodels the soleus transcriptome. (A) Experimental design schematic depicting acute MYC induction protocol and bulk RNA sequencing on MYC-induced (n = 3) and control (n = 3) mice. (B) Principal component analysis of DESeq2-normalized soleus transcriptome. (C) Volcano plot of differentially expressed genes in MYC-induced versus control soleus. Genes meeting thresholds of adj. p < 0.05 and |log2 fold change| > 1 are colored red (upregulated, n = 1,817) or blue (downregulated, n = 1,150); representative genes are labeled. (D-E) Heatmaps of the top 25 upregulated (D) and downregulated (E) differentially expressed genes ranked by adjusted p-value, displaying row Z-scored expression across individual control and MYC-induced samples. (F-G) Gene ontology biological process (GO:BP) gene set enrichment analysis of downregulated (F) and upregulated (G) programs in MYC-induced soleus. Point size reflects gene set size and point color reflects -log10 adjusted *p*-value; pathways with adj. *p*<0.05 are shown.

Differential expression analysis using DESeq2 at an adjusted *p*-value threshold of less than 0.05 identified 2,967 differentially expressed genes (DEGs) in MYC versus control soleus: 1,817 upregulated and 1,150 downregulated (Figure 1C). From the current experiment involving a 48-hour induction of MYC (∼3,000 genes regulated), our prior experiments involving 12 hours of MYC (∼1,500 genes) ^25^, and experiments where MYC was induced constitutively for ≥4 days (>8,000 genes) ^5,22^, we can infer that the extent of altered gene expression pursuant to MYC induction is at least in part dependent on the duration of MYC exposure. Prolonged MYC exposure could directly or indirectly control a much higher percentage of the genome compared to the 15% reported in humans ^38^. The scale of altered gene expression with a 48-hour induction approach is similar to what we find with acute hypertrophic mechanical overload in mice as well as during the 24 hours after a bout of resistance exercise in human skeletal muscle ^39^. Our approach is therefore comparable to exercise according to the overall magnitude of effect on the transcriptome. The 48-hour pulsing strategy aligns with what has been used to induce partial epigenetic reprogramming in skeletal muscle with Yamanaka factors ^27^ and is consistent with our previous approach ^24^.

Gene-level expression following MYC induction is shown in heatmaps of the top 25 up- and down-regulated DEGs ranked by adj. *p* value (Figure 1D&E). There is a consistent effect of MYC that spans ribosome biogenesis factors (*Gnl3*, *Ncl*, *Ftsj3*, *C1qbp*, *Rrp12*, *No1c1*, *Noc21*, *Rpl32*, *Npm1*, *Nop56*, *Wdr77*), RNA modification enzymes (*Ftsj3*, *Mettl1*, *Prmt3*, *Nsun2*, *Nolc1*), telomerase complex components (*Dkc1*, *Nhp2*), and translation-associated genes (*Eif5a*, *Rpl32*, *Cct5*). There is also consistent gene repression across all mice following MYC induction, anchored by muscle identity genes (*Myorg*, *Mybph*, *Itgb1bp2*/*Melusin*, *Tcap*), acetylcholine receptor subunits (*Chrnb1*), and fatty acid and lipid-associated genes (*Slc27a1*, *Acadl*, *Cidea*, *Slc25a20*, *Enho*). We previously showed that downregulation of muscle-specific *Melusin* ^40^ is common to transient MYC induction, Yamanaka factor expression, and endurance+resistance exercise adaptation in murine soleus muscle ^25^. A repression (but not elimination) of certain muscle identity genes is not necessarily deleterious and seems characteristic of the “rejuvenation” signature mediated by exercise-induced MYC in muscle ^25,26^.

A broader gene set enrichment analysis (GSEA) was performed with clusterProfiler against the GO Biological Process database ranked against a background of all 16,105 detected transcripts. The upregulated GSEA landscape (Figure 1F) was dominated by canonical MYC targets such as ribosome biogenesis, rRNA processing, cytoplasmic translation, and ribonucleoprotein complex biogenesis. These data are consistent with the established role of MYC in activating rDNA transcription and ribosome biogenesis in skeletal muscle during hypertrophic loading ^24,41^. Using the same samples used here for RNA-sequencing, we confirm the induction of ribosome biogenesis in the soleus via ∼2-fold upregulation of 45S-ETS, 45S-ITS, and 5.8-ITS after 48 hours of MYC induction ^42^. Enhanced translation via ribosome biogenesis independent from mTORC1 activation, as reported by others in muscle ^28,29^, is the likely predominant contributor to MYC-mediated hypertrophy that we find with repeated transient pulses ^24^.

Beyond the core ribosome biogenesis program, macromolecule methylation and RNA modification (*Mettl1, Nsun2, Trmt61a, Trmt1, Rnmt, Nsun5*) were enriched and may represent a layer of epitranscriptomic and translational control beyond MYC’s influence on overall transcription, as has been found in cancer cells ^43^. Telomere maintenance via telomerase and protein localization to the telomeric region (*Dkc1, Nhp2, Gar1, Nop10, Nat10*) were also enriched, which is consistent with MYC being a known transcriptional activator of telomerase in non-muscle cell types ^44^. The downregulated GSEA landscape (Figure 1G) was organized around three distinct themes. The first was suppression of neuromuscular signaling, with acetylcholine receptor signaling as the most significantly downregulated term (*Chrna1, Chrna2, Chrnb1*). Transient MYC-mediated repression of muscle identity is unsurprising given MYC’s role in epigenetic reprogramming toward pluripotency as a Yamanaka factor ^7,45^. The second theme was repression of immune-related programs and antigen processing and presentation (*H2-Ab1, H2-Aa, H2-Eb1, Tapbp, Psme1*). MYC is a known regulator of tumor immunity regulation and evasion ^46–48^. We now extend these MYC-mediated processes to skeletal muscle. The third theme was a coordinated suppression of lipid catabolic programs (*Slc27a1, Acad1, Fabp3, Acadvl, Hadha, Hadhab, Cd36*), in agreement with our prior work and the work of others ^5,22^ and pointing to an effect of MYC on skeletal muscle metabolism (expanded on below).

### In vitro MYC induction captures in vivo muscle hypertrophic and transcriptomic changes

We speculate that soleus hypertrophy caused by repeated MYC pulses over four weeks we previously observed is due to the cell-autonomous effect of MYC in muscle fibers ^24^; however, indirect (e.g. paracrine) pro-growth effects of muscle-fiber MYC induction on supporting cell types (e.g. immune cells, fibrogenic cells, muscle stem cells) cannot be discounted. MYC transcript is expressed cell-autonomously during myotube contraction and independent from calcium signaling, suggesting its induction is triggered by loading and/or autocrine actions of released cytokines/myokines ^21^. To isolate the cell autonomous effects of MYC induction on muscle fibers, we used a primary cell culture model. Myogenic progenitor cells (MPCs) were isolated from limb muscles of HSA-MYC-GFP mice (n=5 biological replicates) and littermate HSA-GFP mice (no MYC, n=4 biological replicates) using fluorescent activated cell sorting, as previously described by us ^49–52^ (Figure 2A). The sorting strategy uses positive selection for VCAM and a negative selection for non-myogenic cells resulting in high purity myogenic cells ^53^. We have independently validated this antibody-based cell sorting approach using a satellite cell-specific fluorescent reporter mouse model and found high specificity of the antibody to satellite cells using flow cytometry ^50^. After proliferative expansion of MPCs where GFP (and therefore HSA transgene activation) was completely absent, we performed a pilot experiment to determine when the HSA (human skeletal muscle ⍺-actin) promoter becomes active in the presence of doxycycline during primary cell differentiation (Figure 2B). During differentiation with doxycycline in the culture media, GFP (a readout of HSA activation) was scarce at 24 hours when myoblast fusion was not appreciable. By 48 hours, once myoblast fusion had become more pronounced, GFP became more abundant in nascent myotubes. By 72 hours and beyond, when large multinuclear myotubes were present, GFP was enriched in the myotubes (Figure 2B). Thus, HSA does not become active until myotubes begin to form (i.e. upon differentiation). We then assayed for MYC protein after 96 hours of differentiation in doxycycline-treated HSA-MYC-GFP myotubes and found it to be significantly induced, with very low levels in the control condition (doxycycline-treated HSA-GFP MPCs from littermate mice) (Figure 2C). In this model, MYC is present for a similar amount of time as in our *in vivo* approach described above.

**Figure 2.**
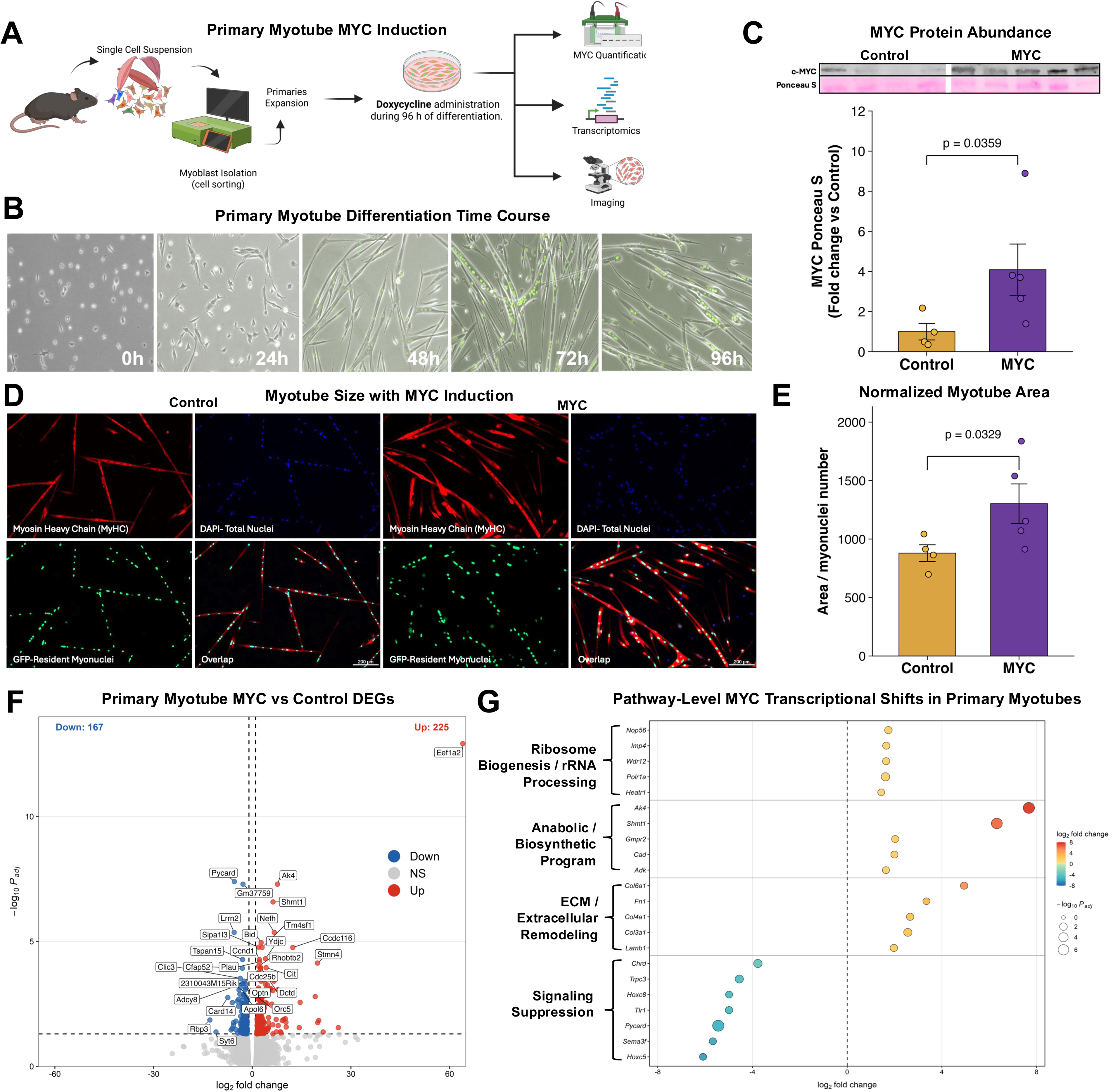
A primary myotube MYC induction model recapitulates *in vivo* growth and transcriptional remodeling. (A) Experimental design schematic for primary myogenic cell isolation and *in vitro* cell culture experiment. (B) GFP fluorescence during primary myotube differentiation, showing progressive activation of the HSA-driven reporter as myoblasts fuse and mature from 24 to 96 hours. (C) Representative western blot and quantification of MYC protein abundance expressed as fold change relative to control. ( D-E) Representative myotube immunofluorescence (D) and quantification of myotube area normalized to myonuclear number (E). (F) Volcano plot of differentially expressed genes in MYC-induced versus control primary myotubes. Genes with Benjamini-Hochberg adjusted *p*<0.05 are colored red for upregulated genes (n = 225) or blue for downregulated genes (n = 167); representative genes are labeled. (G) Curated pathway-level transcriptional shifts in MYC-induced primary myotubes organized into ribosome biogenesis/rRNA processing, anabolic/biosynthetic, ECM/extracellular remodeling, and patterning/signaling suppression modules. Dot size reflects −log10 adjusted p-value and dot color reflects log2 fold change. All experiments performed with biological replicates. Bar graphs show mean ± SEM.

Once establishing our cell culture system as a tractable method for studying MYC in myotubes, we evaluated myotube size in the presence of MYC induction using several biological replicates. Doxycycline was provided in the media for the duration of differentiation (HSA only becomes active upon differentiation in the presence of doxycycline, see above). Consistent with what we observe in muscle fibers *in vivo* ^24^, the induction of MYC causes myotube hypertrophy by 96 hours, measured as MyHC area normalized to GFP+ myonuclei (Figure 2D&E). MYC is known to inhibit differentiation in proliferating myoblasts ^54,55^, again suggesting that MYC’s influence on hypertrophy is attributable to its role in differentiating myotubes following HSA activation. We conclude that MYC can act cell-autonomously to cause myofiber growth and phenotypically reproduces the *in vivo* MYC response ^24^.

Differential expression analysis identified 225 upregulated and 167 downregulated genes (adj. *p*<0.05) (Figure 2F). The most dramatically upregulated gene was *Eef1a2*, encoding eukaryotic elongation factor 1 alpha 2. EEF1A2 is a muscle-enriched translation elongation factor ^56^ whose strong induction is consistent with the capacity for MYC to drive translational elongation machinery. A coordinated multi-gene *Eef* response was identified as common to the human resistance exercise muscle transcriptome and MYC induction in the mouse soleus in our prior work ^24^. Among other significantly upregulated genes were *Ak4*, encoding adenylate kinase 4, which supports mitochondrial energy transfer and nucleotide homeostasis ^57^ and *Shmt1*, encoding serine hydroxymethyltransferase 1, a folate-mediated one-carbon metabolism enzyme that supports nuclear DNA synthesis and DNA methylation. The latter is a recognized MYC transcriptional target in proliferating cells ^58^. Both *Ak4* and *Shmt1* were also significantly upregulated in the acute soleus transcriptome (see Figure 1). The primary myotube model therefore captures a reproducible subset of the MYC-sensitive biosynthetic program observed in muscle tissue *in vivo*.

The curated pathway-level analysis organized the transcriptome response into four modules (Figure 2G). The ribosome biogenesis and rRNA processing module, comprising *Nop56*, *Imp4*, *Wdr12*, *Polr1a*, and *Heatr1*, reflects coordinated induction of nucleolar factors and RNA polymerase I machinery. This set of genes overlaps with the canonical MYC target genes we identified in the *in vivo* acute soleus transcriptome ^24,25^ and aligns with RNA Pol I regulon activation at the onset of loading-induced skeletal muscle hypertrophy ^41^, where *Myc* is strongly induced in myonuclei ^19^. The anabolic and biosynthetic module highlights genes associated with nucleotide metabolism described above, alongside *Gmpr2* and *Srm* which are involved in purine metabolism and polyamine synthesis, respectively. The extracellular matrix (ECM) and extracellular remodeling module contained *Col6a1*, *Fn1*, *Col4a1*, *Col3a1*, and *Lamb1*. MYC induction in differentiating myotubes drives upregulation of basement membrane and interstitial matrix components regulating the ECM. We reported previously on how myofibers can contribute considerably to ECM-related transcription during muscle hypertrophy, including collagen gene expression ^19,59^. The fourth module captured the dominant downregulated effects, with the most strongly suppressed genes including *Lrrn2*, *Pycard*, *Hoxc8* and *Hoxc5*, *Chrd*, *Trpc3*, *Sema3f*, and *Tlr1*. Suppression of *Hoxc8* and *Hoxc5*, which encode HOX-patterning transcription factors often expressed in skeletal muscle precursors, is consistent with MYC-mediated repression of developmental patterning programs in differentiated cells ^60^. Our laboratory ^61–63^ and others ^64,65^ report how changes to *Hox* genes are a prominent feature of exercise adaptation and muscle remodeling across multiple omic layers in mice and humans; again, implicating MYC in the molecular response to exercise ^5,23–26^. Downregulation of *Chrd* and *Sema3f* points to suppression of extracellular morphogenetic and axon guidance signaling. The coordinated repression of *Pycard* and *Tlr1* suggests that MYC induction in primary myotubes suppresses inflammatory and innate immune receptor gene expression.

### Single nucleus RNA-seq (snRNA-seq) after a transient pulse of MYC reveals myonucleus-specific programs related to hypertrophic adaptation and substrate metabolism

To define the potential cell-autonomous effects of pulsatile MYC induction in muscle fibers *in vivo*, we performed single nucleus RNA-sequencing. Soleus muscle nuclei were isolated using fluorescence activated nuclear sorting (FANS) as we have described previously ^19,62,66–69^ (Figure 3A). We pooled tissue from three mice per group following the same 48-hour doxycycline induction and 12-hour washout design in HSA-MYC and control mice (doxycycline-treated littermate HSA only) used in Figure 1. Raw reads were aligned to a custom mouse reference genome that included the human *MYC* transgene sequence ^70,71^, enabling direct detection of transgene-expressing nuclei at single-nucleus resolution. Our data processing workflow is shown in Figure 3B. Because the experiment required pooling of three mice per group to achieve sufficient nuclear yield for library preparation, pseudoreplication was performed prior to pseudobulk differential expression by binning approximately 500 to 1,000 nuclei per bin with a minimum of three bins per condition.

**Figure 3.**
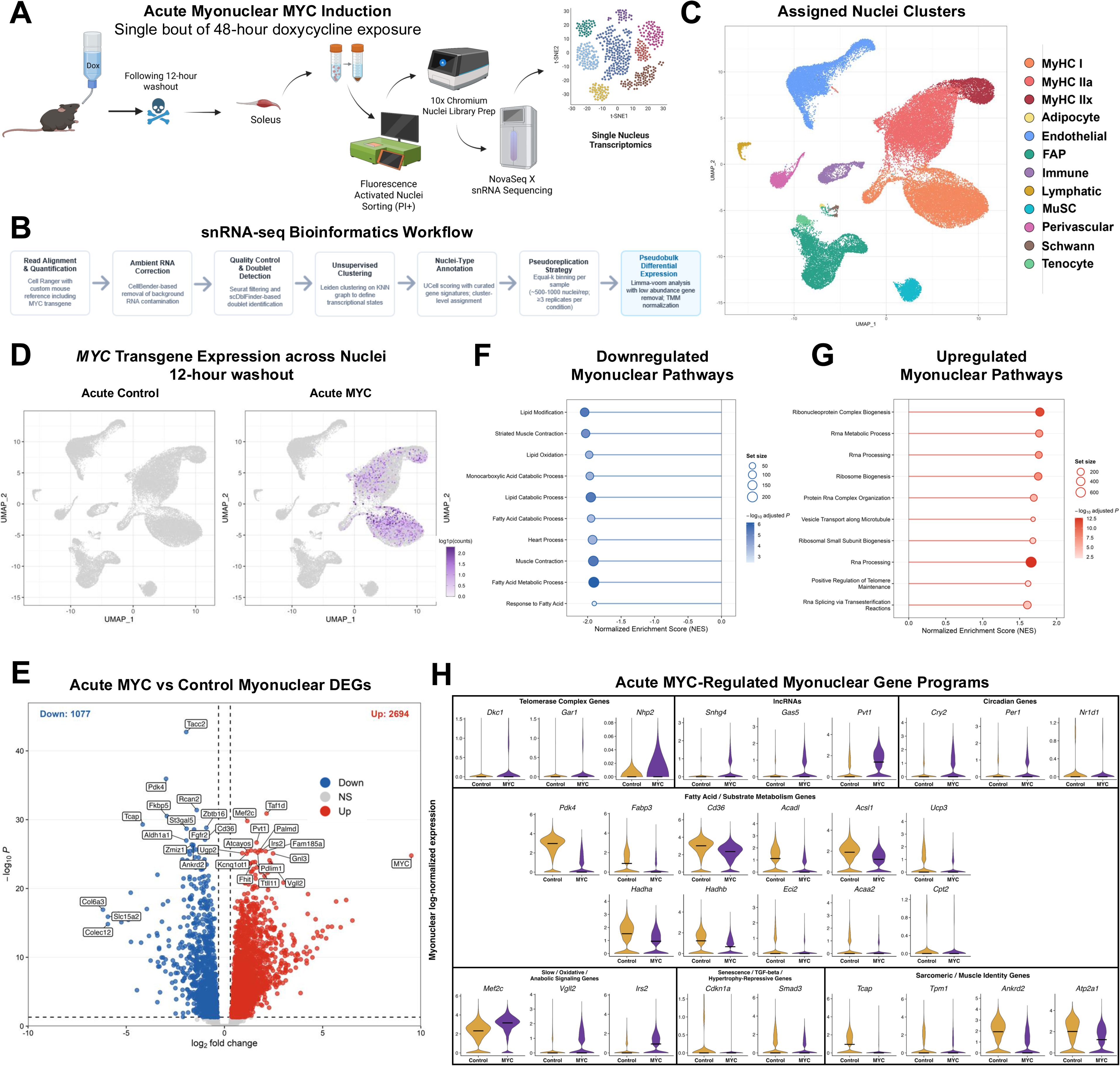
The nucleus type-specific response to a single pulse of MYC in the soleus. (A) Experimental design schematic for acute single-nucleus RNA-seq of soleus muscle following a single 48-hour doxycycline exposure and 12-hour washout in MYC-induced and control mice. (B) Bioinformatic workflow for snRNA-seq processing. (C) UMAP of integrated soleus nuclei colored by annotated nucleus type. (D) Feature plots showing human MYC transgene expression overlaid on the integrated UMAP in acute control, acute MYC 12-hour washout, and acute MYC 24-hour washout conditions. (E) Volcano plot of pseudobulk differential gene expression in myonuclei from acute MYC-induced versus control soleus. Genes meeting thresholds of adjusted *p*<0.05 and |log2 fold change| > 0.30 are colored red for upregulated genes (n = 2,694) or blue for downregulated genes (n = 1,077); representative genes are labeled. (F-G) Top 10 downregulated (F) and upregulated (G) GO Biological Process pathways enriched in acute MYC-induced myonuclei. Pathways are plotted by normalized enrichment score (NES), with point size reflecting gene set size and point color reflecting −log10 adjusted p-value; pathways shown meet adj. *p*<0.05. (H) Violin plots showing myonuclear log-normalized expression of selected MYC-regulated genes grouped into telomerase complex genes, lncRNAs, circadian genes, slow/oxidative/anabolic signaling genes, fatty acid/substrate metabolism genes, senescence/TGFβ/hypertrophy-repressive genes, and sarcomeric/muscle identity genes.

The integrated unified manifold approximation projection (UMAP) resolved eleven major populations, including three myonuclear subtypes (MyHC I [*Myh7* also known as β myosin], MyHC IIa [*Myh2*], and MyHC IIx [*Myh1*]) and nuclei from fibro-adipogenic progenitors (FAPs), endothelial cells, muscle stem cells (muscle satellite cells, MuSCs), immune cells, perivascular cells, tenocytes, lymphatic endothelial cells, Schwann cells, and adipocytes (Figure 3C). MyHC IIb myonuclei were not identified in the dataset, consistent with the well-established near-complete absence of MyHC IIb fibers from the mouse soleus ^72^. Again, adult human limb skeletal muscle only expresses MyHC I, IIa, and IIx isoforms, and not IIb ^35^, similar to the murine soleus muscle. As expected, the largest number of DEGs following MYC induction was in myonuclei.

Plotting the human *MYC* transgene across the acute MYC induction and control conditions showed that transgene expression was almost entirely restricted to myonuclear clusters; 27% of myonuclei were positive for *MYC* transcript compared to near zero detectable expression in other nuclei (Figure 3D; Supplemental Table 1). *MYC* myonuclear transgene expression would intuitively be more abundant across myonuclear types immediately after the cessation of doxycycline, but protein is high at 12 hours after doxycycline cessation in muscle tissue ^24^. This information motivated our tissue collection time point. Our data collectively affirm, at single-nucleus resolution, that transgene induction in the HSA-MYC model is myofiber-specific. Among myonuclear subtypes, MyHC I myonuclei had the highest *MYC* transgene positivity at 34%, compared to 24% in MyHC IIx and 19% in MyHC IIa nuclei. This nucleus-type-specific pattern of MYC expression corresponds to the fiber type-selective hypertrophic response we reported previously with pulsatile MYC induction ^24^. Perhaps preferential Type I fiber hypertrophy is due to greater induction and/or sensitivity to MYC in MyHC I myonuclei. This time course of *MYC* expression in myonuclei relates to the rapid return of MYC protein to baseline within 24 hours of doxycycline removal in muscle tissue established in our previous work ^25^, and validates the 12-hour washout as an appropriate window for capturing the active MYC regulatory state.

Pseudobulk differential expression in myonuclei from the acute MYC versus control comparison revealed a substantial transcriptional response: 2,694 genes were significantly upregulated and 1,077 downregulated at an adjusted *p*-value of less than 0.05 (Figure 3E). Among the most significantly upregulated protein-coding genes after MYC induction were several with well-established roles in nucleolar function and ribosome assembly. *Taf1d,* a TATA-box binding protein-associated factor involved in RNA polymerase I transcription initiation at rDNA promoters, was the most significantly upregulated gene in the dataset. *Gnl3* was highly enriched after MYC induction and is required for ribosome biogenesis and pre-rRNA processing ^73^. *Nop56* and *Nop58* are core components of the box C/D small nucleolar ribonucleoprotein complex involved in rRNA modification and processing ^74^, and *Wdr43* participates in ribosomal small subunit biogenesis ^75^. *Ncl* is a multifunctional nucleolar protein with roles in rDNA transcription and pre-rRNA processing ^76^ and *Mdn1*, an AAA-ATPase essential for pre-60S ribosomal subunit maturation, is a gene we have previously reported to be upregulated with both MYC induction and late-life exercise training in aged mouse soleus ^25^. Altogether, several aspects of rDNA transcription were induced by MYC at the myonuclear level. These gene-level findings are reinforced by GSEA (Figure 3F&G).

MYC is known to activate telomerase ^44^. The telomerase complex genes *Dkc1*, *Gar1*, and *Nhp2* were induced by MYC in myonuclei, consistent with our prior observations that a single MYC pulse upregulates these H/ACA snoRNP components across murine skeletal muscles ^25^ (Figure 3H). The lncRNAs *Snhg4*, *Gas5*, and *Pvt1* were among the more strongly induced non-coding transcripts (Figure 3H). *Gas5* is a growth arrest-specific lncRNA that sequesters glucocorticoid receptor binding activity and is regulated by MYC in other contexts ^77^. *Pvt1* is a well-established MYC-amplified lncRNA target that post-translationally stabilizes MYC protein ^78^. We recently reported that *Pvt1* is upregulated in an *Atf3*+ myonuclear population during acute hypertrophic mechanical overload in adult mice ^66^, and is therefore relevant to skeletal muscle growth. Circadian genes were also strongly regulated by MYC in myonuclei. *Cry2* and *Per1* were induced while *Nr1d1*/*Reverb*⍺ was repressed, indicating that MYC induction in myonuclei influences circadian clock gene balance in myofibers (Figure 3H). Regulation of circadian genes by MYC is consistent with what we observed in plantaris tissue after MYC induction ^19^, as well as the known influence of MYC on circadian output in cancer settings ^79–82^.

The dominant signature for downregulated genes was organized around substrate metabolism, sarcomeric identity, and glucocorticoid-responsive programs. *Pdk4* was the second most significantly suppressed gene in the dataset (Figure 3H). PDK4 phosphorylates and inactivates the pyruvate dehydrogenase (PDH) complex to restrict glucose oxidative flux. *Fabp3*, *Cd36*, *Acadl*, and *Acsl1*, which together mediate aspects of long-chain fatty acid import, intracellular trafficking, and activation for beta-oxidation, were all repressed, as were *Ucp3*, *Hadha*, *Hadhb*, *Eci2*, *Acaa2*, and *Cpt2* (Figure 3H). *Cdkn1a* encoding p21, a well-known marker of cellular senescence, was among the most strongly downregulated genes (Figure 3H). In human skeletal muscle, *p21* (as opposed to *p16*) is the senescence-related marker that is upregulated with aging in myonuclei and causes deleterious outcomes ^83,84^. MYC is implicated in both the progression and elimination of senescence programs in various contexts ^85–90^. MYC modulation of the p21 encoding gene *Cdkn1a* points to the potential for transient MYC to reduce or suppress a senescent-like transcriptome in myofibers. The sarcomeric proteins *Tcap*, *Tpm1*, *Ankrd2*, and *Atp2a1* were also suppressed by MYC (Figure 3H). The downregulation of *Atp2a1* at the myonuclear level by MYC may indicate a downward shift in the fast-type SERCA isoform. *Smad3*, a canonical mediator of TGF-beta and myostatin signaling that limits skeletal muscle hypertrophy, was also significantly repressed, further supporting the interpretation that MYC creates a broad myonuclear program that supports hypertrophy (Figure 3H).

Other genes of interest that are modified by MYC induction in myonuclei include *Mef2c*, a master transcriptional regulator of slow fiber type identity and oxidative gene programs in skeletal muscle ^91–93^ (Figure 3H). The early induction of *Mef2c* following a single MYC pulse may represent a myonuclear ‘priming’ event toward an eventual slower, more oxidative phenotype. *Vgll2*, a co-activator expressed preferentially in slow oxidative muscle that is associated with gene programs for mitochondrial function and contractility in skeletal muscle ^94^, was also strongly induced (Figure 3H). *Irs2*, which encodes insulin receptor substrate 2 and connects insulin and IGF-1 signaling to downstream anabolic pathways, was among the most significantly upregulated signaling genes (Figure 3H). Together, the induction of these transcripts suggests that even a single acute MYC pulse initiates both metabolic and fiber-type-identity remodeling at the level of individual myonuclei.

### A shift in myonuclear proportion and nucleus type-specific metabolic gene expression after repeated bouts of transient MYC induction

Following our “acute” MYC pulse experiment above, we sought to understand how repeated pulses of MYC could alter the molecular landscape in skeletal muscle at single nucleus resolution. We applied the same snRNA-seq approach and workflow outlined above to soleus muscle collected from mice following four weeks of cyclic pulsatile MYC induction (five total pulses, Figure 4A). We used the same 48-hour on, five-day off dosing strategy that we previously showed is sufficient for soleus hypertrophy ^24^. In this experiment, a 24-hour washout period was used to capture the MYC-habituated state, reminiscent of a repeated bout “training” or “lingering” effect. We compared the short-term “MYC trained” response to an acute pulse snRNA-seq experiment that also featured a 24-hour washout to assess the *MYC* transgene (next paragraph). We then compared an acute MYC pulse (12-hour washout) versus the repeated MYC pulse experiment (24-hour washout after the final doxycycline treatment) to understand the effect of a MYC “bout” versus that of MYC “training”.

**Figure 4.**
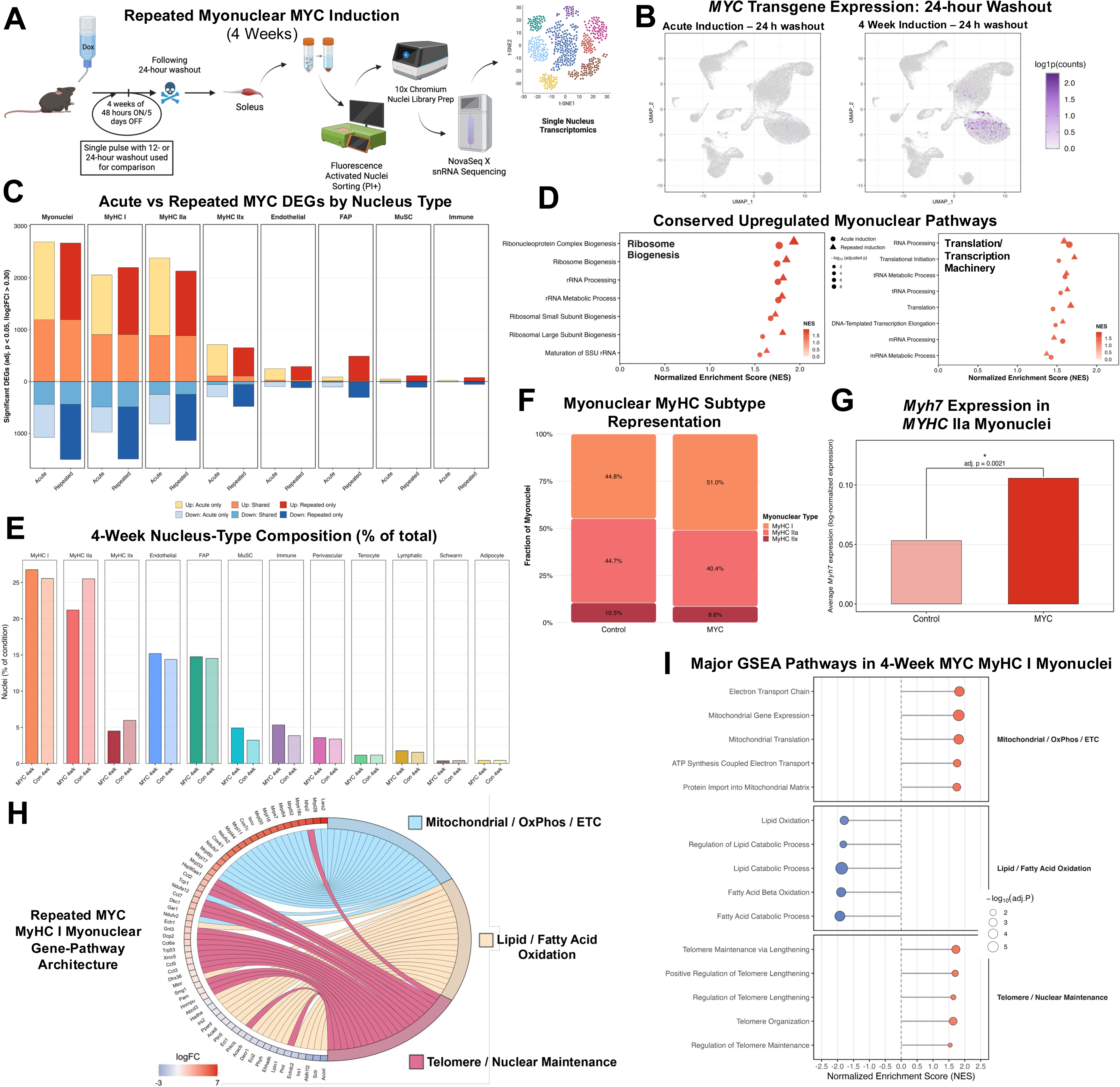
Repeated transient MYC induction over four weeks produces persistent myonuclear transcriptional remodeling and a shift in myonuclear proportion toward MyHC 1. (A) Experimental design schematic for snRNA-seq of soleus muscle following four weeks of repeated pulsatile MYC induction and 24-hour washout in MYC-induced and control mice. (B) UMAP feature plots showing human MYC transgene expression after a 24-hour washout following acute induction and repeated four-week induction. (C) Stacked bar plots showing acute-specific, repeated-specific, and shared differentially expressed genes across myonuclear and non-myonuclear populations. DEGs were defined using adj. *p*<0.05 and |log2 fold change|>0.30. (D) Conserved upregulated myonuclear GO:BP pathways across acute (12-hour washout) and repeated MYC induction, organized into ribosome biogenesis and translation/transcription machinery modules. Circles represent acute induction and triangles represent repeated induction; point color reflects normalized enrichment score (NES) and point size reflects −log10 adjusted p-value. (E) Nucleus-type composition in repeated MYC-induced and control soleus, expressed as the percentage of total nuclei per condition across annotated myonuclear and non-myonuclear populations. (F) Myonuclear subtype composition in repeated MYC-induced and control soleus, expressed as the percentage of total myonuclei assigned to MyHC I, MyHC IIa, and MyHC IIx populations. (G) *Myh7* (MyHC I) expression in MyHC IIa myonuclei from repeated MYC-induced versus control soleus, shown as log-normalized expression. (H) Chord diagram linking repeated MYC-regulated MyHC I myonuclear genes to enriched pathway modules, with gene color indicating differential expression direction and magnitude. (I) Major GO Biological Process pathways enriched in repeated MYC MyHC I myonuclei , organized into mitochondrial oxidative phosphorylation/electron transport, lipid and fatty acid oxidation, and telomere/nuclear maintenance modules. Point size reflects −log10 adjusted p-value, and pathways shown meet adjusted *p* < 0.05.

A notable feature of the repeated pulse dataset was the behavior of the *MYC* transgene following the 24-hour washout period preceding tissue collection. In the acute condition, MYC was not a DEG after a 24-hour washout following 48 hours of induction versus the no-MYC control. Mean normalized transgene expression in myonuclei fell to 0.007 24 hours after an acute MYC pulse, with <1% of myonuclei registering as MYC-positive (Figure 4B). After four weeks of repeated pulsatile induction with the same 24-hour washout, however, MYC remained a significant DEG versus controls. Mean normalized expression in myonuclei was 0.227, with 15% of myonuclei registering as MYC-positive (Figure 4B, Supplemental Table 1). Within the myonuclear subtypes, MyHC I myonuclei retained the highest MYC positivity at 26% compared to 5% in MyHC IIa, while MyHC IIx nuclei showed no detectible MYC signal. Whether this persistence of *MYC* reflects slower transgene mRNA turnover, altered chromatin accessibility at the transgene locus, or changes in the nuclear environment that support sustained transcription following repeated induction remains to be determined. A buildup of doxycycline with repeated pulses is not a likely explanation given the length of washouts after each treatment (five days) and the half-life of doxycycline in mice only being a matter of hours ^95^. Combined with more prevalent *MYC* expression in MyHC I myonuclei, the persistence of *MYC* after repeated pulses may also explain the MyHC I fiber-biased hypertrophy we observe at four weeks ^24^.

Comparing DEGs in the acute (12-hour washout) versus repeated pulse conditions (24-hour washout after final doxycycline treatment) revealed substantial transcriptional remodeling that was both shared and condition-specific across myonuclear populations (Figure 4C). When evaluating all myonuclei irrespective of MyHC type, 1,192 upregulated and 440 downregulated genes were common to both acute and repeated induction. A core transcriptional program is therefore initiated by MYC that is maintained with repeated exposure. Alternatively, 1,478 upregulated and 1,063 downregulated genes were exclusive to the repeated pulse state. This unique pattern was broadly conserved across fiber-type-resolved myonuclear populations: MyHC I myonuclei had 907 upregulated and 490 downregulated DEGs shared with the acute pulse and 1,292 upregulated and 1,004 downregulated genes unique to the repeated pulse condition. MyHC IIa myonuclei had 886 upregulated and 248 downregulated genes shared with acute and 1,246 and 890 repeated-only genes. MyHC IIx myonuclei had the smallest overall transcriptional response: 107 shared upregulated and 59 shared downregulated genes. The primary pathway shared between acute and repeated MYC induction in myonuclei was related to the ribosome and translation, with a stronger effect in the repeated MYC condition (Figure 4D).

Focusing on repeated MYC induction, the nuclear composition of the soleus myonuclear compartment was shifted toward a higher proportion of MyHC I-enriched myonuclei relative to controls (Figure 4E-G). MyHC I-enriched myonuclei comprised 51% of all myonuclei in MYC muscle compared to 45% in controls. MyHC IIx myonuclei represented 9% with MYC versus 11% in controls. The MyHC IIa fraction accounted for the remainder with no MyHC IIb myonuclei represented (Figure 4F). The shift in myonuclear composition toward a slower myosin-type identity is consistent with a progressive remodeling of the myonuclear landscape driven by repeated MYC pulses. This shift is not apparent with an acute MYC pulse. Myonuclear compositional shifts could reflect differential nucleus selection or genuine transcriptional reprogramming of individual myonuclei. Figure 4G provides evidence for the latter by showing *Myh7* (MyHC I) expression specifically within MyHC IIa myonuclei, the myosin class that would be most informative for detecting a genuine “fast-to-slow” transcriptional shift as an “intermediate” myonuclear population. *Myh7* was significantly upregulated in MyHC IIa myonuclei of MYC mice relative to controls (Figure 4G). This shift was accompanied by suppression of *Myh2* in the same population. Even within the MyHC IIa myonuclear compartment, transcriptional identity is biased toward a higher representation of a slower isoform independent from any change in gross nuclear classification according to predominant MyHC type.

We interrogated the repeated pulse MYC state through pseudobulk differential expression and GSEA in MyHC I myonuclei, which had the largest transcriptional response to repeated MYC pulsing. Aside from ribosome pathways (see Figure 4D), three major pathway modules emerged, each represented by the five most significantly enriched GO:BP terms in the analysis (Figures 4H&I). The first and most strongly upregulated module was mitochondrial gene expression and oxidative phosphorylation programs: electron transport chain, mitochondrial gene expression, mitochondrial translation, ATP synthesis coupled electron transport, and protein import into mitochondrial matrix were all significantly enriched. At the gene level, the leading contributors to this module were predominantly mitochondrial ribosomal proteins and oxidative phosphorylation units, including *Mrpl28*, *Lars2*, *Cox7c*, *Cox4i1*, *Iscu*, *Ndufa12*, and *Ndufb3* (Figure 4H). The large effect for mitochondrial ribosomal subunit genes suggests that a consequence of repeated MYC pulsing in MyHC I myonuclei is mitochondrial remodeling. The second module was composed of lipid and fatty acid oxidation pathways that were downregulated in MyHC I myonuclei: fatty acid catabolic process, fatty acid beta-oxidation, lipid catabolic process, lipid oxidation, and regulation of lipid catabolism were all suppressed (Figure 4I). The primary genes anchoring this downregulation included *Irs1*, *Acacb*, *Lpin1*, *Eci1*, *Eci2*, *Decr1*, *Ehhadh*, *Acadl*, *Hadha*, and *Ppard* (Figure 4H). Downregulation of fatty acid genes may represent metabolic reprogramming toward a glucose utilization-biased gene program geared toward biosynthesis and growth ^96,97^. The third module was comprised of upregulated telomere maintenance and nuclear homeostasis in MyHC I-enriched myonuclei (Figure 4H). The genes anchoring this module included the H/ACA snoRNP components *Nhp2* and *Dkc1* alongside *Gar1* and *Gnl3*, which were also among the most significantly induced genes in the acute dataset (see Figure 3). Telomere complex gene induction is a persistent and defining feature of the MYC response in slow myonuclei across both acute and repeated pulse timepoints. The chaperonin complex members *Cct2*, *Cct6a*, *Cct7*, and *Tcp1* were also significant contributors (Figure 4H). *Cct* gene regulation is a shared feature of muscle MYC induction in adult mice and the resistance exercise response in human skeletal muscle ^24^; we now show that this signature is enriched in MyHC I myonuclei. The repeated pulse MyHC I transcriptome also featured significant downregulation of *Dnmt3a*, a DNA methyltransferase. Reduction of *Dnmt3a* in myonuclei is consistent with partial epigenetic reprogramming of the DNA methylome we describe in the context of exercise and MYC induction in the soleus muscle ^24,98^. *Dnmt3a* knockdown and overexpression influences adult skeletal muscle biology ^99,100^ as well as mitochondrial function ^101^. MYC is also known to bind with DNMT3A to repress gene expression ^102^. The functional consequences of *Dnmt3a* suppression specifically in repeatedly MYC-exposed MyHC I myonuclei requires further investigation but may be one additional mechanism through which MYC enhances transcription in skeletal muscle.

### Long-term repeated transient MYC induction causes a faster-to-slower fiber type transition in the soleus

An increase in the proportion of MyHC I-expressing myonuclei revealed by snRNA-seq after four weeks of MYC pulses in the soleus points to a potential fiber type transition from a faster to a slower MyHC phenotype. We report that four weeks of MYC pulses caused muscle fiber hypertrophy but not a fiber type transformation ^24^; however, there was higher variability in fiber type proportion in the MYC pulse group versus controls ^24^. To explore whether MYC can drive a faster-to-slower fiber type transition, we extended MYC pulsing to eight weeks (48 hours on doxycycline, five days off doxycycline for nine total pulses) (Figure 5A). In addition to enhancing soleus muscle mass (Figure 5B, Supplemental Figure 1), extended MYC pulsing caused a significant MyHC IIa-to-MyHC I fiber type transition (Figure 5C and D). The magnitude of this faster-to-slower transition in the soleus is commensurate with what we observe after eight weeks of combined high-volume endurance+resistance progressive weighted wheel running (PoWeR) in the soleus muscles of adult mice ^32,98^, which also induces *Myc* ^98^. MYC therefore represents a single molecular trigger that is sufficient for both muscle hypertrophy as well as fiber type transitioning, driving aspects of a resistance+endurance exercise trained phenotype. Contractile myosin heavy chains are proteins that are unique to striated muscle. Our observation is therefore evidence of a muscle-specific role for MYC that has yet to be observed in the literature. Other hindlimb muscles enriched in MyHC IIb fibers (a feature unique to rodents) and lacking MyHC I, such as the plantaris, do not hypertrophy at the whole muscle level with repeated MYC pulses (Supplemental Figure 2), consistent with our prior report ^24^. The plantaris muscle also experienced a fiber type transition away from MyHC IIa. In contrast to the soleus, the shift was to MyHC IIx - an ultrafast fiber type that does not typically appear in pure form in healthy human muscle ^35,103^. Fiber type transitioning is therefore a conserved feature of MYC pulsing but differs according to muscle type. MyHC IIb fibers are generally much larger than all other fiber types in rodent skeletal muscle ^36^. Plantaris MyHC IIa fibers experienced a modest rightward shift in fiber size but MyHC IIb fibers tended to be smaller (*p*<0.10) after eight weeks of MYC pulses (Supplemental Figure 2). MyHC IIb atrophy could be a sign that IIb fibers are less tolerant than MyHC I and IIa fibers to elevated MYC levels ^5,22^. MyHC IIb fibers are generally less resilient to robust hypertrophic stimuli as they become smaller in the initial phases of muscle mechanical overload ^104,105^ as well as constitutive mTORC1 activation in muscle ^106^.

**Figure 5.**
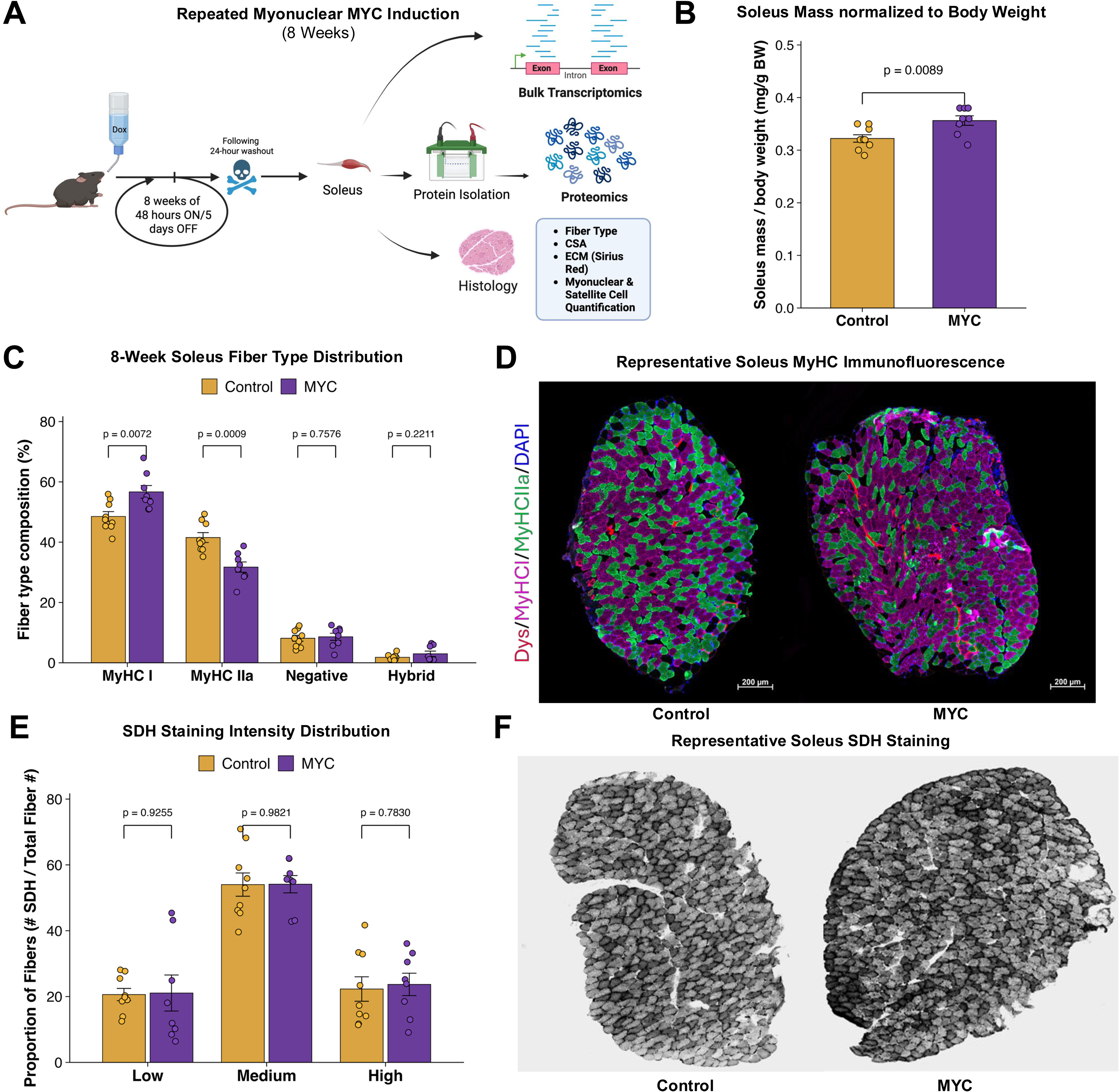
Repeated MYC induces a fast-to-slow fiber type transition in the soleus. (A) Experimental design schematic for 8-week repeated MYC induction and endpoint soleus phenotyping. (B) Soleus mass normalized to body weight in control and MYC-induced mice. (C) Soleus fiber-type composition from MyHC immunofluorescence. (D) Representative soleus cross-sections stained for MyHC isoforms, showing the fiber-type distribution in control and MYC-induced mice. (E) Quantification of SDH staining intensity categories, expressed as the proportion of low-, medium-, and high-SDH fibers relative to total SDH-positive fibers. (F) Representative soleus SDH histochemistry images from control and MYC-induced mice. Bar graphs show mean ± SEM with individual biological replicates overlaid.

In mice, soleus MyHC IIa fibers are more oxidative than MyHC I fibers ^36^. After eight weeks of pulsatile MYC induction, there was no change in the gradient of SDH+ fibers (Figures 5E&F). MYC may at least preserve muscle oxidative potential despite the obvious fiber type shift toward a less oxidative fiber type. Since MyHC fiber type has not shifted significantly by four weeks of MYC pulsing ^24^, but a metabolically reprogrammed signature was clear in myonuclei at this time, we suggest that metabolic changes precede the switch in MyHC type.

Beyond changes in myonuclear proportion revealed by snRNA-seq, the repeated pulse condition produced transcriptional responses in non-myonuclear populations, including fibro-adipogenic progenitors and SCs, that were absent or minimal in the acute dataset (see Figure 4C). Repeated MYC pulses in muscle fibers may therefore have consequences that extend beyond the myonuclear compartment over time. Our results complement those showing that Yamanaka factor induction in muscle fibers remodels the muscle microenvironment to enhance muscle adaptive potential ^27^. MYC induction can reshape the secretome in various cell types ^107–109^. Perhaps MYC in muscle fibers alters cells of the microenvironment via secreted factors, which deserves further exploration. To preliminarily interrogate possible changes in the muscle niche with repeated periodic MYC induction, we performed additional skeletal muscle histology. PAX7+ satellite cell number did not differ between eight-week MYC induced muscle and controls, nor did myonuclear number (Supplemental Figure 3). These data suggest that MYC-mediated hypertrophy may occur independent from satellite cell-mediated myonuclear accretion, which has been reported in a variety of other settings such as mechanical overload, exercise, myostatin depletion, AKT overexpression, and testosterone supplementation ^110–116^. ECM content and muscle lipid content also did not differ between MYC and control conditions (Supplemental Figure 3). This is evidence that there was not pathological disruption to the muscle niche with pulsatile MYC. In response to constitutive MYC induction, muscle fiber damage ensues as evidenced by embryonic myosin heavy chain (eMyHC) expression and IgG infiltration ^22^. We found neither occurrence in our MYC pulsed muscles (Supplemental Figure 3, eMyHC not shown as it was completely absent). Capillary content per fiber was higher with MYC induction concomitant with the fiber type transition (Supplemental Figure 3). MYC drives angiogenesis in tumors via several mechanisms ^117–123^, but how it does so in the context of muscle adaptation is not yet clear and deserving of further investigation.

### Soleus transcriptomics and proteomics shows coordinated responses to MYC pulses are conserved across omic layers and time points

Bulk RNA-seq of the soleus following eight weeks of MYC pulses (24-hour washout after final pulse) revealed 3,172 upregulated and 2,974 downregulated DEGs versus time point-matched controls (Figure 6A); a substantially larger transcriptional signature than the acute response (see Figure 1). We speculate that this is in part due to the persistence of MYC transcript specifically in MyHC I-enriched myonuclei after repeated pulses (see Figure 4), causing an augmented transcriptome profile. While this long-term MYC response comprises fewer altered genes relative to recombination-dependent constitutive MYC induction in skeletal muscle ^22^, ∼6,000 DEGs is still a relatively large number. Nevertheless, this level of MYC induction did not result in a muscle damage transcriptional profile ^124^ that is found with chronic constitutive MYC (i.e. no induction of *Actn2*, *Cdc42*, *Flnc*, *Hspb1*, *Myh10*, or *Dysf*) ^5,22^. We have hypothesized that there is a “goldilocks zone” for MYC induction that is most beneficial for muscle adaptation ^5^. The large amount of transcriptional regulation after eight weeks of pulsing may be approaching the limits of that zone. Perhaps this high level of transcriptomic alteration explains why the hypertrophy we observe in the soleus is not as substantial as we had expected at the muscle fiber level relative to what is observed at four weeks ^24^ (see Supplemental Figure 1). With repeated pulsing of MYC, the early phase of adaptation heavily prioritizes ribosome biogenesis (growth), whereas the latter phases may incorporate mitochondrial adaptations to an increasing extent, resulting in a potential “tradeoff” between adaptive priorities. It is also worth mentioning that high and chronic levels of MYC induction produce a much stronger mitochondrial signature in skeletal muscle ^22^ relative to what is found with acute and transient MYC induction ^5,19,25^. A supraphysiological level of mitochondrial transcriptomic regulation could be a sign of severe metabolic stress ^5,125^.

**Figure 6.**
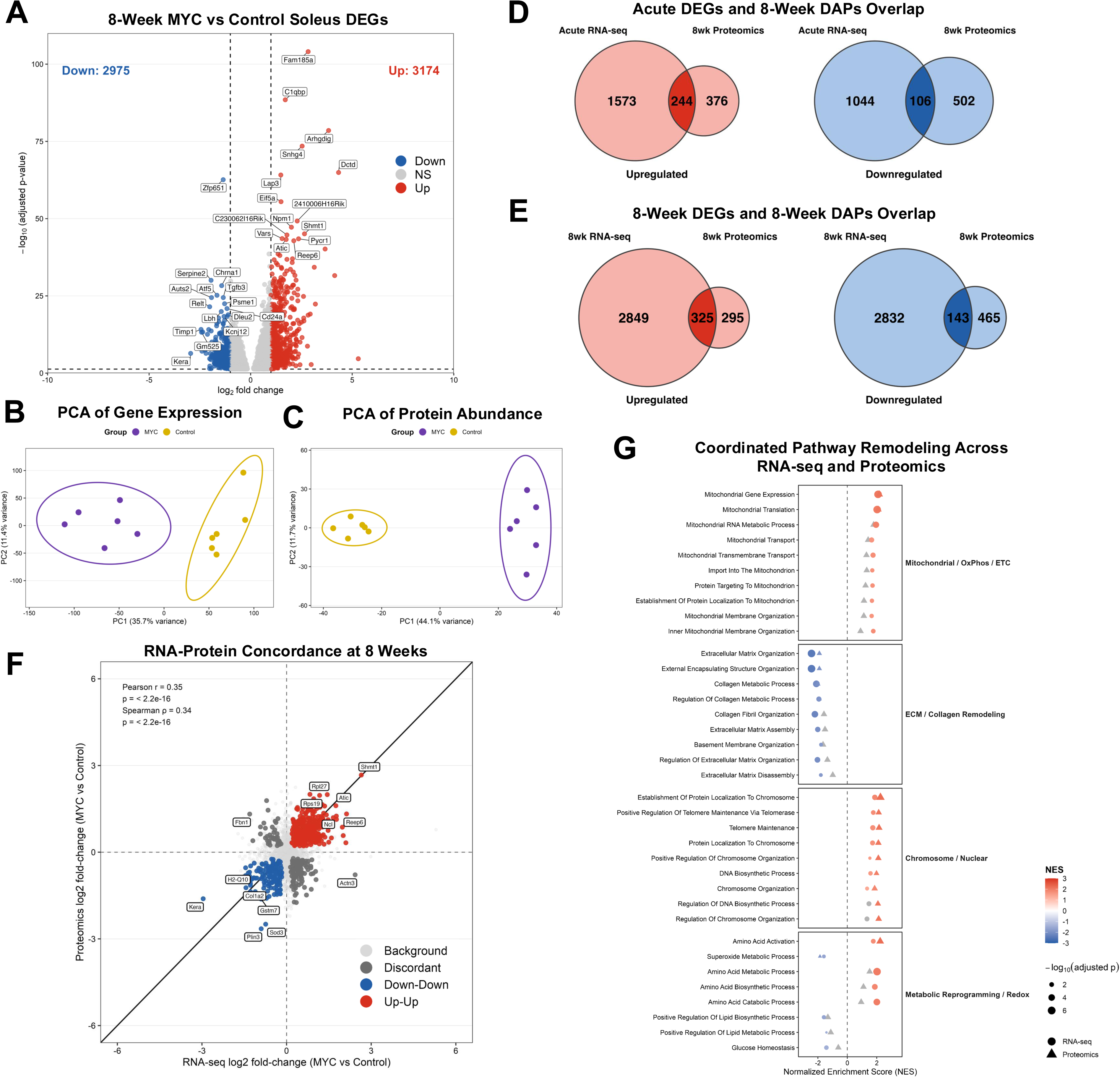
Repeated MYC induction causes coordinated RNA and protein remodeling in 8-week soleus. (A) Volcano plot of bulk RNA-seq differential gene expression in 8-week MYC-induced versus control soleus. Significantly upregulated genes are shown in red (n = 3,171) and significantly downregulated genes are shown in blue (n = 2,972); non-significant genes are shown in gray and representative genes are labeled. (B) Principal component analysis of DESeq2-normalized bulk RNA-seq data from 8-week MYC-induced and control soleus samples, with PC1 and PC2 explaining 35.7% and 11.4% of total variance, respectively. (C) Principal component analysis of mass spectrometry-based protein abundance from 8-week MYC-induced and control soleus samples, with PC1 and PC2 explaining 44.1% and 11.7% of total variance, respectively. (D) Venn diagrams comparing significantly upregulated and downregulated genes from acute soleus RNA-seq with significantly differentially abundant proteins from 8-week soleus proteomics. (E) Venn diagrams comparing significantly upregulated and downregulated genes/proteins from sample matched 8-week soleus RNA-seq and proteomics. Overlap analyses used adjusted *p*<0.05 and |logFC| > 0.1. (F) RNA-protein concordance plot for differentially abundant proteins and genes at 8 weeks, with RNA-seq log2 fold change on the x-axis and proteomics log2 fold change on the y-axis. Points are colored by concordance class: upregulated in both datasets, downregulated in both datasets, discordant, or background. (G) Curated GO Biological Process pathway remodeling across 8-week RNA-seq and proteomics, organized into mitochondrial/OXPHOS/ETC, ECM/collagen remodeling, chromosome/nuclear maintenance, and metabolic reprogramming/redox modules. Circles represent RNA-seq and triangles represent proteomics. Point color reflects normalized enrichment score (NES), point size reflects −log10 adjusted *p*-value, and gray symbols denote pathways that were not significant in that omics layer.

We next sought to understand how eight-week MYC pulse adaptations related to transcriptional and translational profiles in the soleus. Principal component analysis of the transcriptome and proteome from the same samples revealed clear separation between MYC and control samples (Figures 6B&C). We first asked whether the protein-level remodeling evident after eight weeks of repeated transient MYC induction relates to the transcriptional programs that MYC initiates acutely (i.e. a single pulse followed by a 12-hour washout, see Figure 1) (Figure 6D). Of the 620 differentially abundant proteins (DAPs) upregulated by MYC pulses at eight weeks, 244 were also upregulated acutely at the transcriptional level (39%). Of the 608 downregulated DAPs, 106 had correspondingly lower mRNA levels after an acute pulse (17%). We then compared the eight-week MYC pulsed transcriptome to the proteome (Figure 6E). Of the 620 upregulated DAPs, 325 had correspondingly upregulated mRNA levels (52%). Of the 608 downregulated DAPs, 143 had lower mRNA levels (24%). The repeatedly pulsed MYC proteome is therefore more related to the repeat pulse transcriptome than the acute response, especially in the upregulated direction. An RNA-to-Protein fold-change correlation plot quantifies this relationship and shows a positive association with Pearson *r*=0.35 and Spearman rho=0.34 (both *p*<2.2 × 10^−16^, Figure 6F). The moderate but significant correlation means that when a transcript changes significantly at eight weeks, the protein encoding it tends to move in the same direction and with a generally proportional magnitude.

To identify the biological programs shared across omic layers at the eight-week endpoint, we performed GSEA on both the RNA-seq and proteomics ranked gene lists (Figure 6G). The most strongly and consistently enriched upregulated block was mitochondrial gene expression and translation (e.g. MRPL and MRPS proteins, NDUFAF1, NDUFB3, OPA1, TFAM, TOMM40). The second major upregulated set of pathways at the transcriptomic and proteomic levels comprised chromosome and nuclear maintenance functions. The telomere and chromosomal maintenance signature is consistent with the acute response to MYC described above, where *Dkc1*, *Nhp2*, and *Gnl3* were among the most prominently induced transcripts following a single MYC pulse. Four weeks of MYC induction in the MyHC I-enriched myonuclei reveal this same program. The chromosome and nuclear block is anchored by DKC1 and the CCT chaperonin complex, including TCP1, CCT2, CCT3, CCT4, CCT5, CCT6A, CCT7, and CCT8, alongside XRCC5. CCTs are also essential for actin folding ^126^ and are regulated by resistance exercise in human skeletal muscle ^24^. Shared transcriptomic-proteomic downregulated pathways were dominated by extracellular matrix and collagen remodeling programs. The ECM and collagen block is characterized by COL1A1, COL1A2, COL6A1, MFAP4, COMP, LAMB2, CAV1, FBLN5, and FLOT1. This downregulated program contrasts what we observe in our cell culture model where ECM components were upregulated by MYC. The discordance could be related to the *in vitro* environment and myogenic differentiation program, and/or the effects of MYC on non-muscle cells outside of the muscle fiber *in vivo*. The final set of pathways was metabolic reprogramming and redox, which included amino acid activation as a shared program across molecular layers and a shared suppression of superoxide metabolic processes (MAPT, SOD3, PON3, NQO1, NRROS*/*LRRC33, CYGB, GSTP1). Metabolic reprogramming pathways featured SHMT1 among the top-ranked targets. The serine synthesis pathway, via the *SHMT1* gene, was recently implicated in appendicular lean mass in humans ^127^.

### Soleus proteomics shows global maintenance of the mitochondrial proteome in response to long-term repeated MYC pulses and provides evidence of metabolite diversion for hypertrophic and membrane anabolism

With the emergence of the MyHC I fiber-type switch after eight weeks of repeated MYC pulses, we were particularly interested in understanding changes to the muscle mitochondrial proteome (mitoproteome). Using a focused analysis approach, we quantitively evaluated mitochondrial content using the ‘mitochondrial enrichment factor’ (MEF). The MEF is an integrated metric that compares the summed abundance of all mitochondrial proteins (i.e., MitoCarta 3.0 proteins) to total protein abundance determined by proteomics ^128^. This analysis revealed no difference in relative mitochondrial abundance between control and MYC soleus muscles (Figure 7A). We then expanded our analysis to oxidative phosphorylation (OXPHOS) proteins, quantifying the summed abundance of all subunits within each OXPHOS complex ^128^. The relative complex expression between control and MYC soleus muscles was not different for any of the five multi-enzyme complexes of the mitochondrial OXPHOS system (Figure 7B). Consistent with the SDH histology results in the eight-week MYC-pulsed muscles, the proteomics suggest that mitochondrial and OXPHOS content is maintained in response to repeated MYC pulsing, despite the MyHC fiber type shift. Given unique coordinated changes to the mitochondrial transcriptome and mitoproteome as a result of repeated MYC pulsing, however (see above and Figure 4G), these data do not exclude the possibility of changes in mitochondrial efficiency and remodeling. We experimentally explore this possibility below.

**Figure 7.**
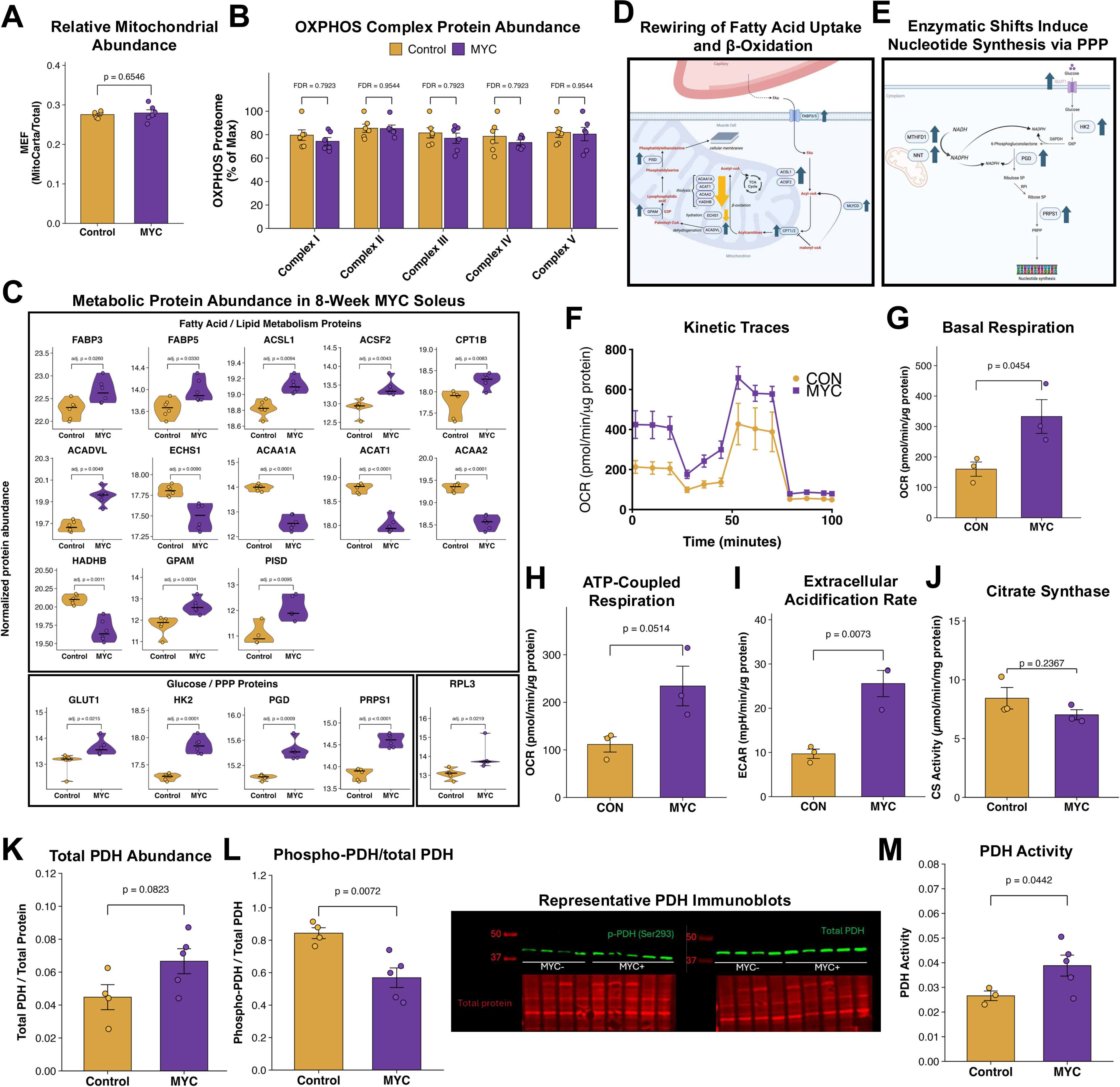
Repeated MYC induction remodels metabolic protein programs toward biosynthetic pathways and enhances oxidative metabolism. (A) Relative mitochondrial protein abundance in 8-week soleus proteomics, expressed as mitochondrial enrichment factor (MEF). (B) Summed abundance of proteins assigned to each oxidative phosphorylation (OXPHOS) complex (Complexes I-V), expressed as percent of the maximal signal within each complex. (C) Curated proteomic violin plots showing normalized abundance of metabolic proteins altered by repeated MYC induction. Adjusted *p*-values from differential abundance analysis are shown above each comparison. (D) Schematic summary of fatty acid uptake, activation, mitochondrial entry, β-oxidation, and phospholipid synthesis pathways altered in eight-week MYC soleus proteomics. Upward and downward arrows indicate the direction of significant protein abundance changes with repeated MYC induction from the proteomics data. (E) Schematic summary of glucose uptake, pentose phosphate pathway activity, NADPH-linked metabolism, and nucleotide synthesis proteins altered in eight-week MYC soleus proteomics. Upward arrows indicate increased protein abundance with repeated MYC induction from the proteomics data. (F-J) Seahorse XF analyses of differentiated control and MYC-induced primary myotubes showing: kinetic data (F), basal respiration (G), ATP-coupled respiration (H), and extracellular acidification rate (I), indicating that MYC increases oxidative and glycolytic metabolism in myotubes. (J) Citrate synthase activity in control and MYC myotubes. (K) Total PDH, (L) Phospho-to-total PDH, (M) and PDH activity in MYC-induced myotubes. Bar graphs represent mean ± SD with individual biological replicates overlaid. Violin plots display normalized protein abundance with individual replicates overlaid and central tendency indicated. Exact *p*-values are shown on the plots; OXPHOS complex comparisons were analyzed by unpaired t-tests with false discovery rate correction, Seahorse summary measurements and citrate synthase activity were analyzed by unpaired t-tests, and proteomic protein-level differences reflect adj. *p*-values from differential abundance analysis.

Similar to the snRNA-seq analyses with repeated MYC pulses, fatty acid metabolism pathways were among significant DAPs after eight weeks of MYC pulses. Proteins responsible for fatty acid cellular entry (FABPs), activation (acyl-CoA ester synthesis, ACSL1, ACSF2), and entry into the mitochondria (CPT1B) were all up-regulated in MYC soleus muscle (Figure 7C). Transient myonuclear transcriptional responses revealed downregulation of similar fatty acid genes and therefore differs from the repeated pulse proteomic response. The disparate signature could be due to a “training effect” of repeated MYC induction and/or be related to the timing of transcriptomic measures (12 hours after MYC induction) versus proteomic measures (24 hours after the final induction).

The enzyme responsible for the first step of fatty acid beta oxidation (dehydrogenation, ACADVL) was upregulated, but enzymes catalyzing the second (hydration, ECHS1) and final steps (thiolysis, ACAA1A, ACAT1, ACAA2, and HADHB) of beta oxidation were downregulated with repeated MYC induction (Figure 7C). Enzymes responsible for the synthesis of phophatidylethanolamine, a vital component of cell membranes from acyl-CoA esters (GPAM and PISD), were up-regulated (Figure 7C). Lipid metabolism is often overlooked in the context of MYC, but is a key feature of MYC-mediated cell growth ^129–132^ as well as somatic cell reprogramming ^133^. Membrane synthesis is a crucial element of cell growth ^134^. As massively voluminous cells, muscle fibers have appreciable membrane surface area that must scale with contractile protein accretion for coordinated and successful MYC-induced hypertrophy. We interpret these findings to mean that pulsatile MYC induction in skeletal muscle may channel fatty acid flux into cell membrane remodeling and/or biogenesis (Figure 7D). Also consistent with an anabolic program, enzymes responsible for cellular glucose entry and metabolism (GLUT1, HK2) and nucleotide synthesis via the pentose-phosphate pathway (PPP) were also upregulated with repeated MYC induction (PGD, PRPS1) (Figure 7C). This signature is indicative of nucleotide synthetic processes that are characteristic of cell growth and are implicated in hypertrophying skeletal muscle ^39,127,135,136^ (Figure 7E). Finally, MYC appreciably increases RPL3 protein (adj. *p* = 0.02) (Figure 7C). The induction of RPL3 is implicated in skeletal muscle growth and plasticity ^137–139^ and may enhance mitochondrial activity in the heart ^140^. MYC regulation of RPL3 is consistent with our prior reports suggesting transient MYC influences this ribosomal protein at the mRNA level across different muscles ^19,25,42^ but is in contrast to what occurs with chronic and constitutive MYC induction that is deleterious to skeletal muscle ^22^.

### Omic responses to MYC induction are corroborated by a coordinated change in oxidative metabolism

We have established that repeated transient MYC pulses are sufficient to induce skeletal muscle hypertrophy *in vivo* and *in vitro*. Our lab and others indicate that muscle hypertrophy with resistance training is in part due to MYC’s role in driving ribosome biogenesis ^19,23,28,29,41,141–143^. The transcriptomic and proteomic data above support this hypothesis. Aside from ribosome biogenesis, however, reprogramming of muscle metabolism by MYC is another prominent feature across the assays we performed, especially following repeated inductions. MYC may affect oxidative metabolism, which is especially compelling based on our snRNA-seq, histochemistry, and proteomic data. Induction of genes such as *Ak4*, *Bcat, Gpt2, Shmt1, Mthfd1l, and Suclg2* also suggest metabolic reprogramming in MYC-induced myotubes. We therefore returned to our *in vitro* platform to determine how MYC may influence metabolism in muscle.

Using a similar experimental approach to what is shown in Figure 2, we induced MYC in differentiating primary myotubes and performed oxygen consumption rate measurements via Seahorse XF Analyzer on myotubes (Figure 7F). MYC was sufficient to promote an appreciably more oxidative muscle phenotype according to basal respiration and ATP-coupled respiration using glucose, pyruvate, and glutamate as substrates (Figures 7G&H). Extracellular acidification rate was also higher with MYC, indicative of augmented glucose usage overall (Figure 7I). In context with the proteomic and histochemical data, MYC induction seemingly improves muscle oxidative potential despite a fiber type transition, combined with muscle growth. The increase in oxidative metabolism could be due to increased mitochondrial efficiency rather than increased mitochondrial content, as citrate synthase enzyme activity (a proxy for mitochondrial mass) did not increase in MYC-induced myotubes (Figure 7J). Total pyruvate dehydrogenase (PDH) abundance was increased, the ratio of phospho-PDH to total PDH (marker of enzyme inactivation) was decreased, and PDH activity was increased in MYC-induced myotubes (Figures 7K-M), collectively supporting increased glucose utilization. There was no difference in *Myhc* gene expression between MYC and control myotubes (all *Myhc*s including developmental and adult, adj. *p*<0.05), again suggesting that a shift in metabolism precedes or is uncoupled from changes in MYHC composition.

### An open access web browser for exploring transient MYC multi-omic data in skeletal muscle

As a resource for the skeletal muscle and MYC biology fields, we provide an open access web browser of all omics data presented in the manuscript: data.myoanalytics.com/study/myc_transient/gene

These data can be queried with a user-friendly interface to promote a deeper understanding of how pulsatile MYC affects skeletal muscle across all the numerous tissue, nucleus, and omic layers we analyzed. This resource will provide new understanding on how an oncogene functions in a highly differentiated cell type and its effect on neighboring non-muscle cells, leading to new insights on cancer resistance as well as exercise adaptation.

## Discussion

In 2012, it was hypothesized that MYC was a central mediator of exercise responses that controlled muscle adaptation, and specifically metabolic adaptations to endurance training ^144^. Since that time, most of the work on MYC in muscle has involved inference and speculation about MYC’s role in loading-induced hypertrophy, and not metabolism *per se* ^19,23,28,29,41,141–143^. This focus seems intuitive since MYC is an oncogene that drives rapid cellular growth in cancer as well as enhanced protein translation via ribosome biogenesis ^2,37,145^; however, MYC-driven growth in cancer is primarily a result of proliferation and not cellular hypertrophy. Skeletal muscle fibers use many of the same anabolic pathways as proliferating cancer cells to grow in response to a hypertrophic stimulus such as resistance exercise ^39^; however, adult muscle fibers typically do not “replicate” to cause muscle fiber growth (e.g. hyperplasia) ^146–148^. We were the first to show evidence that controlled pulses of MYC – mimicking the MYC response to a bout of exercise – was sufficient to induce cellular hypertrophic growth in highly differentiated and non-proliferative muscle fibers *in vivo* ^24^. We further corroborate this finding here *in vitro*. Through our multi-omics explorations, we show that a transient MYC pulse is sufficient to reproduce molecular signatures canonically attributed to exercise adaptation beyond growth, including telomere regulation, extracellular matrix remodeling, and aspects of immunity. We find that processes long described as distinct, parallel adaptations to exercise are each sufficiently influenced by transient MYC. A single pulsatile transcription factor therefore contributes to adaptations previously attributed to multiple independent exercise-sensing pathways. The most striking phenotype we observed was an induction of MyHC I-expressing myonuclei with repeated pulses of MYC. Repeated MYC pulses increased the proportion of myonuclei expressing MyHC I as well as the amount of MyHC I transcript expressed in MyHC IIa-enriched myonuclei. After an eight-week period of MYC pulses, fiber type shifted to a higher proportion of MyHC I fibers at the expense of MyHC II fibers in the soleus. Since myosin heavy chains are not ubiquitously expressed, our work reveals a muscle-specific role for MYC in controlling the proportions of contractile proteins.

Perhaps the most pertinent question based on our findings is: how does MYC control fiber type? We offer several possible explanations. The first is the transcriptional upregulation of *Mef2c* specifically in myonuclei after an acute pulse of MYC. *Mef2c* is a well-documented and powerful regulator of a slow-twitch phenotype in skeletal muscle ^91–93^. After eight weeks of transient MYC pulses, MEF2C was not a significantly upregulated protein. We also cross-referenced our proteomics data against numerous known mechanistic regulators of fast-to-slow fiber type transitioning (including PGC1⍺, Calcineurin, etc.) ^149^ in our proteomics dataset and did not find enrichment of any after long-term repeated MYC induction. Other possible drivers of fiber type transitioning, such as PERM1 ^150^, were downregulated at the protein level with repeated MYC induction. If MEF2C and/or other fiber type-controlling proteins are driving a MYC-mediated induction in MyHC I fibers, it may be a rapid and transient event that cannot be captured by a proteomic snapshot at our chosen time point, or the protein abundance was too low to be detected. Another potential explanation for how MYC may control fiber type transitioning is via changes that are secondary to alterations at the neuromuscular junction (NMJ). In the acute RNA-seq data, acetylcholine receptor signaling was the most significantly downregulated term. While speculative, MYC may induce NMJ remodeling which could possibly contribute to fiber type transitioning ^151–153^. A third explanation is that MYC directly controls MyHC genes by binding MyHC promoters to drive expression or unbinding from promoters to lower expression. MyHC IIa, IIx, and IIb genes all contain E-boxes ^154^ - the known binding motif for MYC - whereas MyHC I has MCAT sites ^155^. MyHC I expression may be mediated by *cis* mechanisms such as enhancers ^156,157^, epigenetic modifications ^158^, g-quadruplexes ^159^, and/or post-translationally ^160,161^. A fourth possibility is that all events are occurring simultaneously. Future investigations will explore these possibilities.

Recent evidence suggests myosin heavy chains can direct muscle metabolism ^162^, implying contractile identity leads metabolism. Our data suggest the opposite: that metabolic changes precede the change in fiber type. A MYC-mediated myonuclear signature of oxidative metabolism genes appears prior to the fiber type transition detected by histology ^24^. We observe signs of metabolic reprogramming towards biosynthesis, which is associated with muscle hypertrophy in mice and humans ^127,135,163,164^. We find an induction of HK1, HK2 and SLC2A1 (GLUT1) in the soleus proteome pointing to increased glucose handling potential in the presence of MYC, as well as increased PPP/nucleotide synthesis enzymes; however, there are also indicators of enhanced oxidative metabolic potential across omic layers and time points that can be traced to changes in myonuclear transcription. We corroborated a MYC-mediated increase in oxidative metabolism using oxygen consumption rate analysis in MYC-induced myotubes and PDH activity assays. In addition to ribosome biogenesis, MYC is strongly implicated in mitochondrial biogenesis ^165–168^. We observe an appreciable 1.3-fold induction of MT-CO1 (Complex IV subunit, 1.3 LogFold, adj. *p*=0.0001) in our proteomics data. MT-CO1 tightly correlates with mitochondrial translation and biogenesis from endurance training in mice ^169^. Furthermore, MYC induction in fibroblasts agrees with our *in vitro* mitochondrial respiration experiments concomitant with increased mitochondrial mass ^166^; however, our *in vitro* data and *in vivo* proteomics both point to mitochondrial remodeling versus mitochondrial biogenesis with MYC induction in skeletal muscle. In the adult murine heart, MYC induction alters mitochondrial and fatty acid oxidation markers, all seemingly independent from PGC1⍺ induction, which collectively contributes to cardiac stress resilience ^167^. Altogether, it appears that transient MYC induction results in both greater aerobic glucose metabolism for energy production and greater anaerobic glucose metabolism for biomass production in skeletal muscle. Combined with signs of a shift in fatty acid prioritization toward membrane synthesis, our data establish a unified metabolic framework that involves several biosynthetic pathways to support muscle adaptation via transient MYC. Future investigations involving flux analyses and tracers can validate metabolic shifts and the potential shunting of fatty acids towards membrane synthesis, as well as more granular measures to determine mitochondrial volume, density, and size distribution using high-resolution microscopy. We conclude that MYC, which is transiently but strongly induced by exercise in human skeletal muscle, mediates hypertrophic, metabolic, and perhaps stress-resilience benefits associated with chronic exercise training.

### Limitations of the Study

It will be important to identify the ideal “dose” of MYC and to mechanistically tease out the most deterministic features downstream of MYC activation that mediate hypertrophic, metabolic, and MyHC fiber type adaptations. This information could aid in the development of skeletal muscle therapies that do not carry the risk of off-target oncogenesis. From a functional standpoint, testing whether transient MYC-induced muscles are more adaptive to endurance or resistance exercise training, have enhanced performance, are resistant to atrophy, can combat nutrient stress, and/or are refractory to aging across biological sexes ^5,26^ will be revealing. While questions about MYC in muscle remain, our work implicates MYC as a central regulatory node in exercise-induced skeletal muscle adaptation ^170^.

## Resource Availability

Raw and processed RNA-sequencing data from all bulk and single-nucleus experiments reported in this study will be deposited in the Gene Expression Omnibus (GEO). All analyzed data are found here: data.myoanalytics.com/study/myc_transient/gene

## Acknowledgements

This study was supported by NIH grants AG063994, AG080047, and AG088465 to K.A.M. This work was performed while KAM was a Glenn Foundation for Medical Research/American Federation for Aging Research Junior Investigator Awardee. Proteomics was funded by a voucher from the IDeA National Proteomics Resource (R24GM137786) to RGJ. SE was supported by the Swedish Society for Medical Research postdoctoral grant (PG-24-0429), the Swedish Research Council for Sports (P2025-0154 & P2026-0203), and the Wenner-Gren Foundations. FvW was supported by the Swedish Research Council (no. 2022-01392) and the Swedish Research Council for Sport Science (P2024-0102; P2025-0171). BFM was supported by a VA Research Career Scientist award (IK6BX007150). Thank you to Tyrone A. Washington, PhD for providing tissue from high fat-fed mice as a control for one of the experiments. The TRE-MYC mice were provided as a generous give from Andrew P. McMahon, PhD.

## Author Contributions

Conceptualization: RGJ, SE, FvW, KAM

Data curation: RGJ, NS, TC, RW, MET, JEM, AKB, MT, JAEM, JLF, AI, YW

Formal analysis: RGJ, NS, TC, AKB, MT, RW, JAEM, JLF, AI, KAM

Funding acquisition: KAM

Investigation: RGJ, PJK, NS, ARC, FM, TLC, MET, AKB, MT, RW, JAEM, JLF, ALZ

Methodology: RGJ, ARC, JLF, CSF, BFM, AI, YW, KAM

Project administration: KAM, JLF, BFM, YW

Resources: SE, FvW, CSF, JLF, BFM, AI, YW, KAM

Software: RGJ, AI, YW

Supervision: KAM

Validation: RGJ, YW, KAM

Visualization: RGJ, ARC, AI, KAM

Writing – original draft: RGJ, AI, YW, KAM

Writing – review & editing: RGJ, PJK, NS, ARC, FM, TLC, MET, AKB, MT, RW, JAEM, ALZ, SE, FvW, CSF, JLF, BFM, AI, YW, KAM

## Declaration of generative AI and AI-assisted technologies

AI was not used for the development of any aspect of this work.

## Supplemental Information Titles and Legends

**Supplemental Figure 1. Muscle mass and fiber cross sectional area (CSA) in the soleus muscle after eight weeks of repeated transient MYC pulses.** Absolute muscle mass (A) and fiber CSA (B) in MyHC I, MyHC IIa, MyHC-negative (IIx) fibers from eight-week control and MYC-induced soleus muscle. Bars show mean ± SEM with individual biological replicates overlaid. \**p*<0.05, ^#^*p*<0.10.

**Supplemental Figure 2. Muscle masses of hindlimb muscles and plantaris muscle histology after eight weeks of repeated transient MYC pulses.** Quantification of absolute (A) and normalized mass (B) across hindlimb muscles after eight weeks of transient MYC pulses. Plantaris fiber type distribution (C) and muscle fiber size distribution (D). Bars show mean ± SEM with individual biological replicates overlaid.

**Supplemental Figure 3. Satellite cell abundance, myonuclear number, ECM content, immune infiltration, and capillarity after 8 weeks of transient MYC pulses in the soleus.** (A) PAX7+ nuclei frequency in 8-week control and MYC-induced soleus muscle. (B) Myonuclear number in 8-week control and MYC-induced soleus muscle. (C) Sirius Red-positive ECM content in 8-week control and MYC-induced soleus muscle. (D) Representative images or quantification of lipid accumulation in 8-week control and MYC-induced soleus muscle. (E) Representative IgG staining assessing immune/injury-associated infiltration in 8-week control and MYC-induced soleus muscle. (F) Quantification of capillaries per myofiber in 8-week control and MYC-induced soleus muscle. MYC induction increased capillaries per myofiber (unpaired two-tailed t-test, *p* = 0.0019). Bar graphs show mean ± SEM with individual biological replicates overlaid.

## Methods

### Ethical Approval and Animal Housing

All animal procedures were approved by the University of Arkansas IACUC (AUP 21037 and AUP 24032). Mice were housed in a temperature and humidity-controlled room maintained on a 12:12-h light-dark cycle with food and water provided *ad libitum* throughout experimentation. Animals were euthanized via cervical dislocation following deep anesthesia induced by inhaled isoflurane. All mice for *in vivo* experiments were females >4 months of age but <10 months of age.

### Skeletal Muscle Fiber-Specific MYC Induction Model

The doxycycline-inducible, skeletal muscle fiber-specific MYC induction mouse model (Human skeletal muscle ⍺-actin reverse tetracycline transactivator tetracycline response element [Tet-On] MYC, or HSA-MYC) used throughout this study was generated as previously described by us ^19,24,25^ (Jackson Laboratory Strains 038301 and 19736). All mice were heterozygous for each transgene, and doxycycline was used as the tetracycline analog to induce muscle fiber-specific MYC expression *in vivo*. Littermate HSA-only mice treated with doxycycline served as controls for all *in vivo* experiments. A subset of HSA-MYC mice was crossed with a tetracycline response element green fluorescent protein mouse (Jackson Laboratory Strain 005104) to generate HSA-MYC-GFP mice, which were used for primary cell culture experiments with littermate HSA-GFP as controls.

### Acute MYC Induction

HSA-MYC and littermate control mice (n = 3 females per group) were given doxycycline in their normal drinking water with sucrose (0.5 mg·mL^-1^ with 2% sucrose) for 48 hours at 6-8 months of age. Doxycycline-supplemented water was then replaced with normal drinking water in the morning, and mice were euthanized 12 hours after the removal of doxycycline unless otherwise specified (a subset were analyzed after a 24-hour washout for snRNA-seq).

### Repeated Pulsatile MYC Induction

To explore the outcome of long-term repeated MYC induction in skeletal muscle, we repeated (four week) and extended the repeated pulsatile induction design (eight week) first described in our prior work ^24^. Adult female HSA-MYC and littermate control mice (n = 8 and n = 9 respectively) were given doxycycline in their normal drinking water with sucrose (0.5 mg·mL^-1^ with 2% sucrose) for 48 hours, followed by five days of normal drinking water. This pulse approach was repeated for a total of five or nine inductions over four or eight weeks. All mice were euthanized 24 hours following the final doxycycline treatment, in the morning before 11:00 AM (a subset were analyzed after a 12-hour washout for snRNA-seq).

### Primary Myoblast Isolation then Differentiation with MYC Induction

Primary myogenic progenitor cells were isolated from several juvenile (9–13 weeks of age) male HSA-MYC-GFP and HSA-GFP mice using a protocol adapted from the Rando laboratory ^53^. Briefly, limb muscles were dissected, minced, and enzymatically dissociated using collagenase and dispase, and filtered through a 40 µm strainer to generate a single-cell suspension. The cell suspension was then incubated at 4°C with antibodies against VCAM, CD31, CD45, and SCA1, on ice and a biotinylated secondary antibody was used for VCAM. Cells were pelleted (500 x g for 5 minutes), re-suspended, and sorted by fluorescence-activated cell sorting (FACS) on the MACSQuant Tyto Cell Sorter (Miltenyi Biotec, Bergisch Gladbach, Germany). Primary myogenic cells were identified and sorted based on viability detected by an absence of propidium iodine (PI) (non-viable cells) and negative for CD31/CD45/SCA1 (FITC, “dump” channel). The sorted positive selection was for VCAM-1 high cells (PE-Cy7). Debris were excluded using back/side scatter profiles.

Sorted cells were plated on culture dishes coated with 1% extracellular matrix (ECM) gel (Sigma-Aldrich E1270; derived from Engelbreth-Holm-Swarm murine sarcoma) diluted in serum-free DMEM (Gibco 11965-092) supplemented with 1% penicillin-streptomycin (Corning 30-002-CI). Cells were expanded in growth medium, which was replaced daily, consisting of Ham’s F-10 medium (Gibco 11550-043), 10% horse serum (Gibco 16050-122), and 1% penicillin-streptomycin, supplemented with 25 μg/mL fibroblast growth factor (FGF; Sigma F0291) prepared in 0.1% bovine serum albumin (BSA). Cells from passages 2–4 were used for protein analysis, RNA extraction, and myotube area normalized to myonuclear number. Cells were plated at 700 cells/cm² density and allowed to reach ∼65% confluence before differentiation. Differentiation was induced by replacing growth medium with differentiation medium consisting of DMEM, 5% horse serum, and 1% penicillin-streptomycin, and 2 μg/mL doxycycline. Cells were maintained under differentiating conditions for 72 or 96 hours.

For protein abundance, cells were harvested in 2X sample buffer and assessed by Western blotting using an anti c-MYC primary antibody (Cell Signaling Technology, D84C12, 1:500). To measure myotube area, cells were fixed in 4% paraformaldehyde and stained for myosin heavy chain (MyHC; DHSB, MF20, 1:100 in 2% BSA/PBS). Myotube area was quantified from fluorescence images acquired at a constant intensity in the AF555 channel using a Zeiss microscope and normalized to the number of GFP-positive myonuclei. Statistical comparisons for MYC protein abundance and myotube area per myonuclear number were performed using one-tailed Welch’s t-tests, n = 4–5 biological replicates per group, with values presented as mean ± SEM.

### Bulk RNA Isolation, Sequencing, and Analysis

TRIzol Reagent (Sigma-Aldrich, St. Louis, MO, USA) was used to isolate RNA from primary myotubes and for soleus muscle. For tissue, RNA was isolated from approximately half of each soleus (4–6 mg) for the acute induction experiment and from 2–4 mg of soleus tissue for the 8-week endpoint experiment using. Myotubes and tissue were homogenized using zirconia beads and a Fisher Bead Mill (Fisher, Hampton, NH, USA). Following homogenization, RNA was isolated via phase separation by addition of chloroform then centrifugation. The aqueous phase was transferred to a new sterile tube and further processed on spin columns using the Direct-zol Kit (Zymo Research, Irvine, CA, USA). For the 8-week soleus samples, poly-acryl carrier was added to improve RNA yield from lower input tissue mass (Molecular Research Center, Cincinnati, Ohio, USA). Concentration and purity were determined using a BioTek Take3 micro-volume microplate with a BioTek PowerWave XS microplate reader (BioTek Instruments Inc., Winooski, VT, US).

Isolated RNA from all samples was sequenced by Novogene (Sacramento, CA, USA) on an Illumina HiSeq platform using 150 bp paired-end sequencing. Raw FASTQ files were processed in Partek Flow. Reads were aligned to the mouse reference genome (GRCm39) using STAR 2.7.8a and quantified against the mm39 annotation model. Features with a maximum count < 5 across all samples were removed prior to downstream analysis for noise reduction. Normalization and differential expression analysis were performed using DESeq2, with differentially expressed genes (DEGs) identified at a Benjamini-Hochberg false discovery rate adjusted *p*-value < 0.05 with no fold-change cutoff was applied for DEG classification. Gene set enrichment analysis (GSEA) was performed in R using the clusterProfiler package ^171^ against the Gene Ontology Biological Process database. All expressed genes passing the low-expression filter were included in the ranked input to avoid false pathway enrichment. Gene sets with a Benjamini-Hochberg adjusted *p*-value of < 0.05 are reported.

### Fluorescence-Activated Nuclear Sorting and Single-Nucleus RNA Sequencing

Soleus muscle nuclei were isolated *via* Fluorescent Activated Nuclear Sorting (FANS) on a MACSQuant Tyto Cell Sorter (Miltenyi Biotec, Bergisch Gladbach, Germany). Soleus muscle from three mice per condition were pooled in 1mL of homogenization buffer (500 µL HEPES [1 M], 3 mL KCl [1 M], 250 µL spermidine [100 mM], 750 µL spermine tetrahydrochloride [10 mM], 10 mL EDTA [10 mM], 250 µL EGTA [100 mM], 2.5 mL MgCl [100 mM], 5.13 g sucrose) with 4 µL of Propidium Iodide (PI), and 0.2U/µL RNAse inhibitors (Protector RNase Inhibitor, Millipore Sigma, Burlington, MA, USA). The muscle was initially minced in buffer with scissors on ice in a low-bind 1.5 mL tube until a slurry, dounced ∼20 times with a plastic pestle with a gentle twist at the bottom, then strained through a 20-μm MACSQuant pre-separation filter (Cat #: 130-101-812) directly into MACSQuant Tyto high-speed sorting cartridge (Cat #: 130-121-549). FANS gating was established to exclude debris and identify nuclei positive for PI (intrinsic DNA label), then sorted directly into PBS/1% BSA/RNase inhibitors to minimize dilution. Nuclei suspension concentration was measured using the automated Countess II (ThermoFisher Scientific, Waltham, MA, USA), aiming to load ∼20,000 nuclei within the optimal range for the 10X chromium chip. Master mix (MM) solution was combined with up to 36.6 µl of the nuclei suspension + PBS, then loaded into the GEM-X 3’ chip for GEM formation. Library was prepared using the Single Cell 3’ Reagent Kit v4 according to the manufacturer’s protocol. Following library construction, libraries were sequenced on the Illumina Nova NextSeq X Plus System by Novogene to 200 million reads per library.

### Single-Nucleus RNA Sequencing Analysis

Raw FASTQ files were aligned and quantified using Cell Ranger v9.0.0 (10X Genomics) against a custom mouse reference genome (GRCm39) augmented with the full coding sequence of the human MYC transgene. Inclusion of the transgene in the reference was necessary to enable direct detection and quantification of transgene-expressing nuclei at single-nucleus resolution, consistent with the myofiber-specific MYC overexpression design of the experiment. Following alignment, gene-barcode matrices were subjected to ambient RNA correction using CellBender (v0.2.0) ^172^, which applies a deep generative model to distinguish true biological transcripts from background contamination.

Corrected .h5 feature files were imported into R (v4.5) and processed using Seurat v5. Nuclei were filtered to remove low-quality barcodes based on total UMI counts, detected gene counts, and mitochondrial transcript fraction, with a mitochondrial read cutoff of < 5%. Doublets were identified and removed using scDblFinder. Data were normalized using log-normalization, and highly variable features were identified for dimensionality reduction. Principal component analysis was performed, and the number of PCs retained for downstream graph construction was selected using an automated elbow and cumulative variance criterion, with a minimum of 15 PCs and an 85% cumulative variance target. A shared nearest-neighbor graph was constructed using k = 30.

Unsupervised clustering was performed using the Leiden algorithm implemented via igraph v2.2.2 ^173^. The Leiden algorithm was selected over the more commonly used Louvain algorithm because it provides explicit guarantees of well-connected communities and prevents the formation of disconnected or weakly connected partitions. Multiple resolutions were evaluated, and the final resolution of 0.4 was selected from candidates with modularity within 95% of the maximum observed modularity, prioritizing highest mean silhouette width in PCA space and then fewer clusters. Cell type identity was assigned using a two-step consensus annotation workflow. UCell (v2.14) ^174^ enrichment scores were computed from normalized RNA expression using curated marker gene panels canonical to the various nuclei types in skeletal muscle. These marker-based scores were then reconciled with cluster topology and used to label cluster identity.

Because the experiment required pooling of nuclei from three biological replicates per group to achieve sufficient nuclear yield for the library preparation described above, direct single-nucleus differential expression testing would treat individual nuclei as independent observations, inflating type I error rates ^175^. To address this, pseudoreplicates were generated using a dynamic binning strategy that aggregated nuclei within each biological sample. For each nuclei type and condition, nuclei were partitioned into bins targeting approximately 500 to 1,000 nuclei per replicate as larger binning experiences diminishing returns to the strength of the differential expression testing ^175^, with a minimum of three pseudoreplicates per group required for inclusion in differential expression analysis.

Pseudobulk count matrices were converted to edgeR DGEList objects (edgeR v4.4.2) and normalized using TMM with low-abundance genes removed using filterByExpr. Count data were transformed using the voom method and fitted with limma (v3.62.2) linear models. Differential expression was tested using limma:: treat at a primary absolute log2 fold-change threshold of 0.30, with a secondary high-confidence threshold of 0.585 also applied. Genes were considered significant at Benjamini-Hochberg FDR < 0.05. Gene set enrichment analysis was performed using fgsea (v1.32.4) with mouse-native MSigDB C5 Gene Ontology Biological Process gene sets obtained via msigdbr (v26.1.0). Genes were ranked by moderated t-statistic, and enrichment was performed using fgseaMultilevel. All expressed genes passing the low-abundance filter within each nuclei type stratum served as the background gene list for each comparison, consistent with other pathway analyses in this study. Enriched pathways were considered significant at FDR < 0.05 and were separated into positive and negative normalized enrichment score categories. All analyses were performed in R (v4.5) with Bioconductor (v3.21).

### Histology

Soleus muscles were embedded in optimal cutting temperature compound, frozen in liquid nitrogen-cooled isopentane, and stored at -80°C until sectioning. Transverse cryosections of 8 µm thickness were cut using a cryostat and air dried for a minimum of one hour prior to staining. Fiber cross-sectional area and fiber type distribution were assessed by immunohistochemistry as previously described ^32,105^. Soleus sections were incubated in primaries for MyHC I (BA-D5, DSHB), MyHC IIa (SC-71, DSHB), dystrophin primaries. Plantaris sections were stained using MyHC IIa, MyHC IIb (BF-F3, DSHB), and dystrophin then the appropriate isotype-specific for MyHC I, MyHC IIa, MyHC IIb) before applying DAPI. All images were captured using an upright fluorescent microscope at 20X magnification (Zeiss Axiolmager M2, Oberkochen, Germany) where whole muscle sections were imaged using the mosaic function in Zeiss Zen 3.8.3 for Microsoft.

For succinate dehydrogenase (SDH) enzyme histochemistry, 7 µm sections from the soleus muscle were air dried and incubated in a succinate buffer containing 50mM Sodium Phosphate pH at 7.6, 50mM sodium succinate, and 0.5mg/ml nitroblue tetrazolium for 45 minutes at 37°C. After incubation, tissues were rinsed in deionized water and then processed for fiber typing using the same protocol previously described. Analysis was conducted using MyoVision v2.1 which simultaneously assesses fiber type and SDH intensities using bright field and fluorescent microscopy images taken at the same time. Pixel intensities for each fiber were exported and assessed using 8-bit grey scale from 0-255 where black pixels are given a value of zero (arbitrary units) representing darker SDH staining and lighter pixels are given higher values representing weaker staining. For binning of SDH staining, arbitrary thresholds of 0-100 for dark staining, 101-175 medium staining, and greater than 175 for light staining of fibers were applied. Fibers were counted and summed across full soleus cross-sections and assessed for group differences.

Sirius Red staining was performed to evaluate extracellular matrix organization and collagen deposition as described ^113,176^. 7 µm soleus sections were incubated in Bouin’s fixative (Biolyst inc., 15990) for 1 hour at 56°C and rinsed with deionized water for 2 minutes followed by a 2-hour incubation period in PicroSirius Red solution (Biolyst inc., 26357-02) and washed in 0.5% acetic acid, dehydrated in 95% and 100% ethanol, and equilibrated in xylene for ∼5 minutes before mounting with cytoseal (Biolyst Inc.,18009). Regions of interest were then imaged using an inverted Nikon (Eclipse Ti) microscope at 10x. Images were then analyzed using ImageJ software for percentage of red area.

Satellite cell abundance was assessed by staining for PAX7+ nuclei as previously described ^113^. Briefly, soleus tissue sections were fixed with 4% PFA for 7 minutes and washed for 3 rounds of 3 minutes in PBS, treated with 3% hydrogen peroxide to block endogenous peroxides for 7 minutes, followed by an epitope retrieval step in sodium citrate (2.94 g/L pH 6.8) at 92°C for 11 minutes and allowed to cool to room temperature and washed for 3 rounds of 3 minutes in PBS. Sections were then blocked with mouse-on-mouse IgG blocking reagent (Vector, #MKB-2213) for 45 minutes followed by 3 rounds of PBS washes, and then with 1% TSA with 0.1% Triton at room temperature for 1 hour. Anti-Pax7 (ab_528428, DSHB) and laminin (Sigma L9393) primary antibodies were then applied overnight at 4°C. Following biotinylated secondary antibody (1:1000) treatment in 1% TSA (Jackson Immunoresearch, 115-065-205) for 70 minutes and 3 rounds of 3 minutes PBS washes, tissues were then treated with SA-HRP (1:500; Thermo Fisher, S-911) and laminin (anti-rabbit AF647; 1:200) secondaries for 1 hour at room temperature before another round of PBS washes and applying the TSA AF594 amplification for PAX7 for 15 minutes. Nuclei were stained with Prolong Diamond anti-fade mountant with DAPI (Invitrogen, P36962). Nuclei that were both PAX7+, DAPI+ and resided withing the laminin boarder were manually counted as satellite cells with relative counts calculated as PAX7+ nuclei per fiber where fibers were assessed using the semi - automatic image analysis tool MyoVision ^177,178^.

IgG staining was performed as a negative control to confirm the absence of myofiber membrane damage as described by Ham et al. ^22^. Briefly, 7 µm sections from the soleus were cut and allowed to air dry before incubating in cold acetone fixing for 10 minutes followed by 3 washes for 3 minutes each in PBS. Sections were then blocked in 5% BSA for 20 minutes followed by an IgG anti-rat secondary antibody (Jacksonimmuno, 112-165-143 ) in 1% BSA over night at 4°C.

For lipidspot (Biotium Inc., cat #70065) staining, ∼7 µm sections were cut and stained according to the manufacture’s recommendations. Briefly, the tissues were removed from -80°C storage and allowed to dry for at least 1 hour before fixing in 4% PFA. The tissues were then washed in PBS and incubated in a PBS and triton-X (0.5%) solution for 20 minutes. The same primary antibody for dystrophin was applied as the above IHC stains overnight at 4°C. Tissues were then rinsed in PBS before the dystrophin secondary antibody was applied for 1 hour at room temperature. Before the lipidspot incubation there was another round of PBS rinses. The lipidspot stain was diluted in 1:100 in PBS and 0.1% triton-X and applied to the tissues for 30 minutes, rinsed in PBS, and mounted with a glycerol-based media before imaging. For a positive control, we used an aged (∼22 month) high fat fed mouse ^179^ (60% of kcal from fat) tibialis anterior muscle that was cut and stained at the same time as the HSA-MYC and HSA-rtTA (no MYC control) tissues.

For immunohistochemistry (IHC) analysis of capillarity, PECAM-1 (Cat#: 2H8-supernatant) monoclonal antibody developed and deposited to the DSHB by Bogen, S.A., created by the NICHD of the NIH and maintained at The University of Iowa, Department of Biology, Iowa City, IA 52242 was used for determination of soleus capillarization markers (capillaries per mm^2^, capillaries per myofiber, and capillaries in contact with each fiber (CCEF)). Muscle sections air dried for 15 mins then fixed in cold acetone (−20°C) for 10 mins followed by 2x5 min washes in PBS. Sections were then blocked for 30 min in 5% normal horse serum (NHS) in PBS. Primary antibodies for PECAM-1 (1:5) and dystrophin (1:250, PA1-37587, Invitrogen, Carlsbad, CA, USA) in PBS with 5% NHS were completed for 1 hour at room temperature then overnight at 4°C. The following day sections were washed 3x5 min in PBS and then incubated in secondary antibodies against PECAM-1 (1:500, AF594, goat anti-Armenian hamster IgG, Invitrogen) and dystrophin (1:200, AF350, donkey anti-rabbit IgG, Invitrogen) for 1 hour at room temperature. Following secondary incubation, sections were washed 3x5 mins with PBS and mounted with 50/50 glycerol and PBS solution for imaging. Images were captured at 20x total magnification at room temperature with a ZEISS upright microscope (Axio Imager M2, Oberkochen Germany). For analysis of capillarization, each sample had multiple ROIs (225x225µm) captured (2-4 per sample) for analysis of capillary density. Image analysis was performed by the same investigator who was blinded to sample condition throughout the analysis using Zeiss ZEN microscopy software. Capillaries per myofiber were calculated only with completely visible myofibers (i.e. myofibers cut off at the edges were not included in analysis) within each ROI. All capillaries that were in contact with the complete myofibers were counted and then divided by the total myofiber count in each ROI. Unpaired t-test with test for normality and significance value set at p<0.05 were used to determine differences in soleus muscle capillarization with and without 8 weeks of muscle-specific MYC induction.

### Proteomics Sample Preparation

Proteomics was performed on soleus tissue from the same animal used for 8-week bulk RNA-seq in each biological replicate, with approximately two-thirds of each soleus allocated to proteomics and the remaining one-third used for RNA isolation, enabling a within-animal matched multi-omic design. Frozen soleus tissue was cryopowdered and homogenized in 1X RIPA buffer. Protein concentration was determined using Pierce 660 protein quantification assay (Thermo Scientific). Sixty micrograms of total protein per sample was desalted by acetone precipitation overnight at -20°C. The protein pellet was resolubilized in Laemmli buffer (Bio-Rad) with 50mm DTT and 20 µg of protein per sample was resolved by SDS-PAGE. Gels were fixed and stained with GelCode Blue Stain Reagent (Thermo Scientific). To reduce the dominance of high-abundance structural proteins inherent to skeletal muscle, including titin and myosin heavy chain, the lower portion of each gel lane was excised for downstream analysis. A standard in-gel digestion method was used. The gel pieces were washed to remove the GelCode Blue Stain and then reduced in 10 mg/mL DTT, alkylated in 35 mg/mL iodoacetamide, dried, and then rehydrated and digested overnight with 1 μg trypsin per sample in 200 μL 50 mM ammonium bicarbonate. The mixture of peptides was extracted from the gel, evaporated to dryness in a SpeedVac, stored at -20 °C until resuspended for liquid chromatography (LC)-tandem mass spectrometry (MS) analysis.

### Proteomics Analysis

Mass spectrometry acquisition and protein identification were performed by the UAMS National Proteomics Core. Proteins were identified by searching against the mouse UniProt/Swiss-Prot reference proteome. Protein intensities were normalized using variance stabilizing normalization (VSN). Differential abundance analysis was performed using a limma-based linear modeling framework with Benjamini-Hochberg false discovery rate correction. Proteins with an adjusted p-value < 0.05 were considered significantly differentially abundant, without imposition of a hard fold-change cutoff, consistent with the approach described above for bulk RNA-seq differential expression analysis in this study.

All downstream visualization and pathway analyses were performed in R (v4.5). Gene set enrichment analysis of differentially abundant proteins was performed using fgsea (v1.32.4) with mouse-native MSigDB C5 Gene Ontology Biological Process gene sets obtained via msigdbr (v26.1.0). Proteins were ranked by moderated t-statistics and enrichment was performed using fgseaMultilevel. Enriched pathways were considered significant at FDR < 0.05. Mitochondrial protein content was quantified using the mitochondrial enrichment factor (MEF), calculated by comparing the summed abundance of all MitoCarta 3.0-annotated proteins to total protein abundance ^128^. Oxidative phosphorylation complex abundance was assessed by quantifying the summed abundance of all detected subunits within each of the five OXPHOS complexes ^128^.

### Mitochondrial Stress Test and Citrate Synthase Activity

Cellular respirometry of myotube cells was measured using the Seahorse Extracellular Flux Analyzer (Agilent Technologies) as previously described ^180^. Briefly, control and MYC myoblast cells were seeded in extracellular matrix-coated XFe24 cell culture microplates in growth medium at a density of 6x10^4^ cells per well, as described above. Following a 24-hour incubation, growth medium was replaced with differentiation medium and doxycycline, and cells were allowed to differentiate for 72 hours as described above. On the day of the assay, cells were washed once with prewarmed Seahorse XF DMEM assay medium supplemented with 1 mM pyruvate, 2 mM glutamine and 25 mM glucose, pH 7.4 (Agilent Technologies), then incubated in assay medium in a non-CO_2_ incubator at 37°C for 1 hour. The mitochondrial stress test was conducted using consecutive injections of 2.5 mM oligomycin (Cayman Chemical), 2 mM carbonyl cyanide-p-trifluoromethoxyphenylhydrazone (Cayman Chemical), and 0.5 mM rotenone/0.5 mM antimycin A (Cayman Chemical, Ann Arbor, MI) as the final concentrations. Measurements were normalized to protein concentrations as determined by the BCA protein assay (Thermo Fisher Scientific).

Citrate Synthase (CS) Activity Assay (Cayman Chemical) was performed to assess mitochondrial mass in CON and MYC cells following 72 hours of differentiation (as described above). Briefly, cells were washed with PBS, detached from the plate with a cell lifter, and pelleted by centrifugation. Pellets were resuspended in the CS assay buffer, sonicated for 3 seconds and incubated on ice for 10 minutes. Cell suspensions were centrifuged at 10,000 *x g* for 5 minutes at 4°C to remove insoluble material. Cellular extracts were diluted 1:50 prior to the assay in CS assay buffer and the assay was performed and quantitated according to the manufacturer’s instructions. CS activity was normalized to total protein concentrations as determined by the BCA protein assay (Thermo Fisher Scientific).

### Western Blotting

For Western blotting, samples were separated by SDS-PAGE using 4-15% Criterion TGX Gels (Bio-Rad). Total protein stains were visualized using a Revert Total Protein Stain Kit (LI-COR Biosciences) for total protein normalization of target bands. Total PDH was assessed using a PDHE1a antibody (1:1000, NBP2-33922, Novus Biologicals), and phosphorylated PDH was assessed using an anti-phospho-PDHE1a (Ser293) (1:1000, NB110-93479, Novus Biologicals). IRDye fluorescent secondary antibodies were used, and membranes were visualized using an Odyssey Imaging System (LI-COR Biosciences). Target band densitometry was quantified using Image Studio Software (LI-COR Biosciences).

PDH activity was assessed using a microplate assay kit (ab109902, Abcam). Briefly, wells of the microplate were coated with an anti-PDH monoclonal antibody to capture fully-intact, functionally-active PDH enzyme complexes. PDH activity was determined as the reduction of NAD+ to NADH, coupled to the reduction of a reporter dye and monitored as the absorbance at OD450 nm for 30 minutes, using a Biotek Synergy H1 Multimode reader. Equal amounts of protein from each sample were added to wells in duplicate, and data are presented as the change in absorbance over time.

### Statistical Analyses

Statistical analyses were performed in R and GraphPad Prism, as appropriate for each assay. For non-omic two-group comparisons, data were analyzed using unpaired t-tests unless otherwise specified. Welch’s correction was applied when variance assumptions were not met. Two-tailed tests were used unless a directional one-tailed test was prespecified, as in the primary myotube MYC protein abundance and myotube area per myonuclear number analyses. For assays involving multiple related endpoint comparisons, including OXPHOS complex abundance and fiber type or staining-bin comparisons, individual group comparisons were performed with multiple-testing correction where indicated. Significance for non-omic assays was defined as *p* < 0.05. Data are presented as mean ± SEM unless otherwise stated. Bulk RNA-seq differential expression was performed using DESeq2, with Benjamini-Hochberg adjusted *p* < 0.05 used to define differentially expressed genes. Single-nucleus pseudobulk differential expression was performed using edgeR/limma-voom with limma::treat, using a primary absolute log2 fold-change threshold of 0.30 and Benjamini-Hochberg FDR < 0.05. Proteomic differential abundance analysis was performed using a limma-based linear modeling framework following variance-stabilizing normalization, with Benjamini-Hochberg FDR < 0.05 defining differentially abundant proteins. Gene set enrichment analyses were performed using preranked approaches as described above, and pathways were considered significant at FDR < 0.05.

