## Supplementary figures and images for "Transient MYC Mimicking the Exercise Response Orchestrates Multifaceted Skeletal Muscle Adaptations"

### Supplemental Figures

#### Supplementary Figure 1

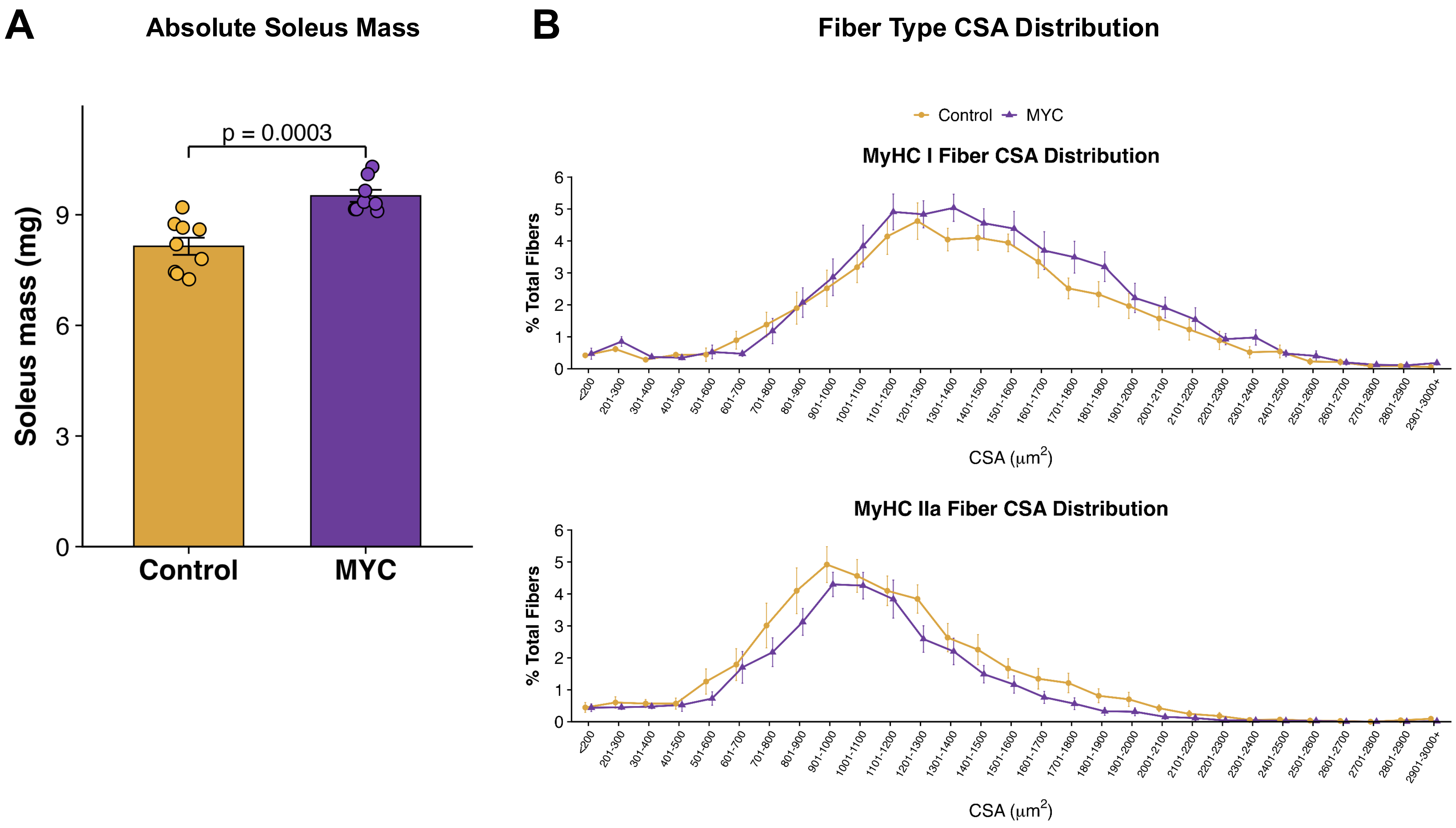

#### Supplementary Figure 2

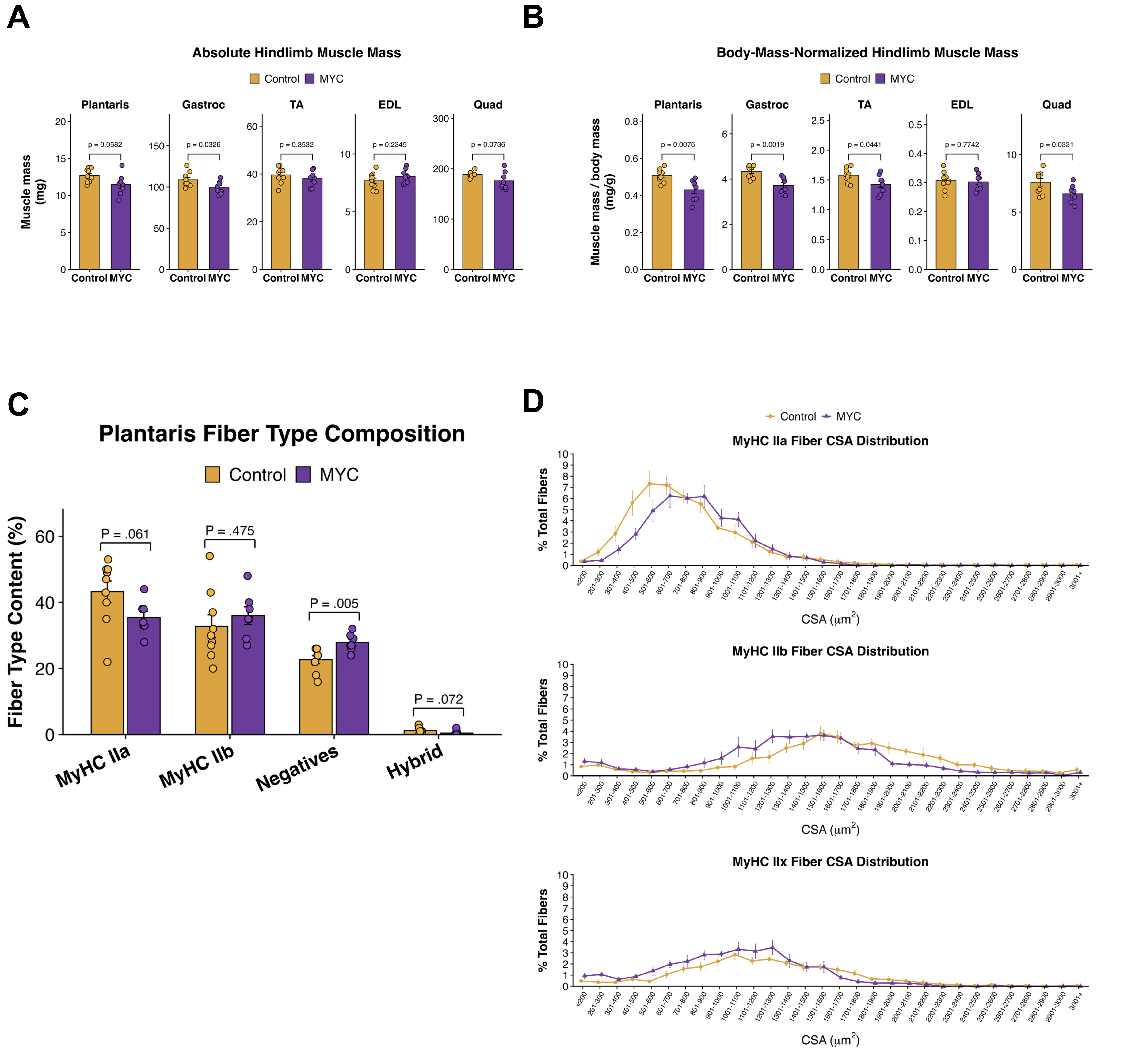

#### Supplementary Figure 3

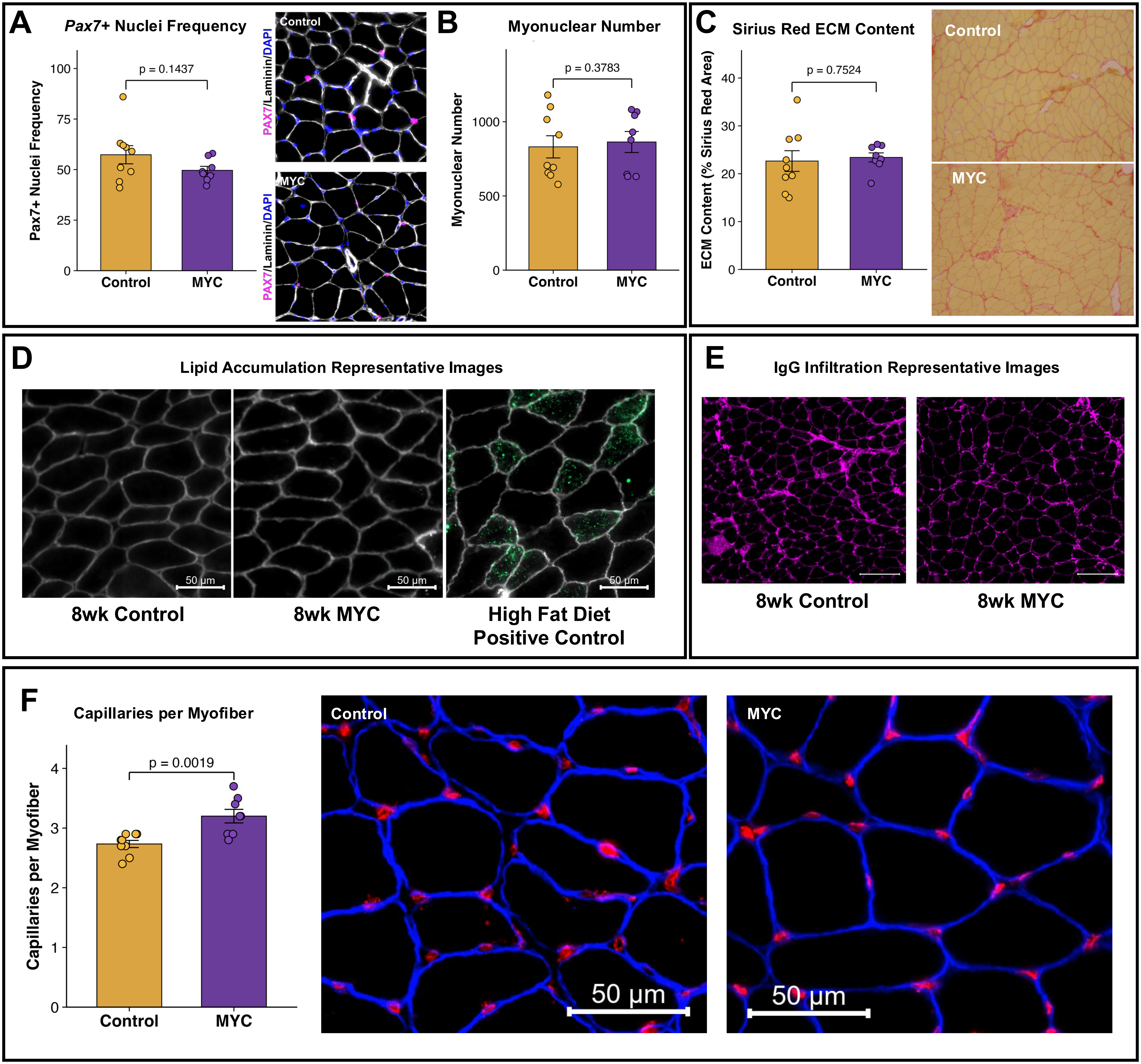
